# Conservation applications of niche modeling: native and naturalized ferns may compete for limited Hawaiian dryland habitat

**DOI:** 10.1101/2023.11.01.565183

**Authors:** Krystalyn Edwards-Calma, Laura Jiménez, Rosana Zenil-Ferguson, Karolina Heyduk, Miles K. Thomas, Carrie M. Tribble

## Abstract

**Premise:** Competition from naturalized species and habitat loss are common threats to native biodiversity and may act synergistically to increase competition for decreasing habitat availability. We use Hawaiian dryland ferns as a model for the interactions between land-use change and competition from naturalized species in determining habitat availability.

**Methods:** We use fine-resolution climatic variables and carefully curated occurrence data from herbaria and community science repositories to estimate the distributions of Hawaiian dryland ferns. We quantify the degree to which naturalized ferns tend to occupy native species suitable area and map remaining available habitat given land-use change.

**Results:** Of all native species, *Doryopteris angelica* has the lowest percentage of occurrences of naturalized species in its suitable area while *D. decora* has the highest. However, all *Doryopteris* spp. had a higher percentage overlap—while *Pellaea ternifolia* had a lower percentage overlap—than expected by chance. *D. decora* and *D. decipiens* have the lowest proportions (*<* 20%) of suitable area covering native habitat.

**Discussion:** Areas characterized by shared environmental preferences of native and naturalized ferns may also decrease due to human development and fallowed agricultural lands. Our study demonstrates the value of placed-based application of a recently developed correlative ecological niche modeling approach for conservation risk assessment in a rapidly changing and urbanized island ecosystem.

## INTRODUCTION

One of the primary goals of conservation biology is to assess the amount and location of suitable habitat for species of concern (e.g., Stage 2 in Smith et al., 2018). Correlative ecological niche models (hereafter referred to as CENMs) use occurrence data (records of where individuals of the species have been observed in the past) to estimate the climatically suitable conditions of a particular species (Busby, 1991; Carpenter et al., 1993; Hirzel et al., 2002; Phillips et al., 2006; Soberón, 2010; Stockwell, 1999). CENMs may allow conservation practitioners to determine the extent of suitable habitat for threatened species, prioritize areas for land conservation, identify regions where a rare species may exist but still be undetected, and many more possible applications (Engler et al., 2004; Ferraz et al., 2021; Guisan et al., 2013; Larson et al., 2004; Villero et al., 2017; Zhu et al., 2013). However, often applications of correlative ecological niche modeling confuse theoretical and practical differences between the fundamental and realized niche, confounding the interpretation of such analyses and thus the proper application of CENMs in conservation (Peterson and Soberón, 2012; Soberón, 2010; Zhu et al., 2013).

The fundamental niche is defined as the set of environmental conditions that a species could inhabit in the absence of biotic factors (such as competition) and dispersal limitations, while the realized niche is the set of conditions that a species currently inhabits as a result of the dynamics of biotic interactions, abiotic factors, and historical dispersal patters (Hutchinson, 1957; Pulliam, 2000). For the vast majority of niche modeling studies, occurrence data from biological collections describe where a particular species has been observed and represent a sample from the realized niche, which may under-represent the fundamental niche of a species (Jiménez et al., 2019; Soberón and Arroyo-Peña, 2017). The dearth of appropriate statistical methods to estimate the fundamental niche from presence-only data had limited the ability of conservationists to expand their efforts to preserve areas where vulnerable species can live—or historically lived—because the right climatic conditions exist, even if the vulnerable taxa currently do not occur in a given location due to biotic interactions. Therefore, it is beneficial to infer fundamental niches for conservation purposes.

While direct measurements of the absolute abiotic limits of organismal function and survival is an ideal way to estimate the fundamental niche, these data may be unavailable for rare species of conservation concern and may be time consuming and expensive to collect, thus statistical approximations of the fundamental niche are often more feasible for many conservation applications. New developments in correlative ecological niche modelling provide statistical approximations of the fundamental niche and thus may allow conservation practitioners to preserve critical areas where species can survive. These new CENM modeling approaches are improving our ability to infer the fundamental niche by assuming a hypothetical shape for the response function of a species to the environment where fitness declines proportionally with distance from centroid according to a multivariate normal model (which produces an ellipsoidal shape, see Jiménez et al., 2019), and by combining presence data with a second source of information, such as the accessible area that the species is able to explore (see Jiménez and Soberón, 2022).

Additionally, factors beyond the abiotic variables used in most CENMs may significantly affect the suitable area for species of conservation concern. For example, land-use change may mean that areas characterized by the appropriate climatic conditions for a species to thrive are instead unavailable due to land conversion from native habitat to areas of intense human development, agriculture, or invasive-dominant ecosystems. Additionally, competition with naturalized species with similar habitat preferences may also significantly affect the amount of available area for native species of conservation concern. These factors are typically not accounted for in traditional CENMs (but see Ashraf et al. (2021); Gutiérrez et al. (2014); Peterson et al. (2006); Sallam et al. (2013)), so conservation applications of CENMs may overestimate the available suitable area of species of interest. In this study, we describe a pipeline for modeling the fundamental niche using the statistical approach in Jiménez and Soberón (2022), while accounting for the impacts of land use change and competition from naturalized species on available suitable habitat. We apply this methodology to the vulnerable native Hawaiian dryland ferns in the family Pteridaceae and discuss the potential future applications of such approaches for conservation.

Hawai‘i is widely known as a biodiversity hotspot, with textbook examples of adaptive radiations following long-distance dispersal to the isolated volcanic archipelago (e.g., the lobeliods, Givnish et al. 2009, silverswords, Baldwin and Sanderson 1998; Witter and Carr 1988, and songbirds, Lovette et al. 2002). Despite the small area of available land, Hawai‘i contains an extraordinary diversity of biomes and environmental conditions, which has likely allowed organisms to adapt to novel environments and diversify (Barton et al., 2021; Gagne and Cuddihy, 1990; Kitayama, 1996; Price and Wagner, 2004; Vitousek, 2004). Moreover, because of Hawai‘i’s isolation and the rarity of long distance dispersal events, rates of endemism are particularly high: 90% of the approximately 1300 native Hawaiian vascular plant species are found nowhere else (Baldwin and Wagner, 2010; Rønsted et al., 2022; Wagner et al., 1999). Sadly, Hawai‘i’s spectacular biodiversity is also under immense threat (Sakai et al., 2002). A recent report on the conservation status of approximately 1000 native Hawaiian vascular plants showed that 75% of the flora is threatened or endangered, while only 33% of that flora receives protection under the U.S. Endangered Species Act and the Hawai‘i State Government, implying that much of the native flora is under threat yet unprotected (Rønsted et al., 2022).

Loss and endangerment of tropical flora is particularly striking in tropical dry forests, which is one of the most threatened biomes worldwide (Cordell and Sandquist, 2008; Quesada et al., 2009; Wilson et al., 1988). They once made up 42% of the world’s tropical regions but currently only 2% of those forests remain (Buzzard et al., 2016). Across the tropics, the dry forest biome has attractive features for human use: they tend to be suitable areas for livestock and agriculture because their soils are often fertile and they possess a marked rainfall season, and they are favored for human development because the dry environment may reduce the risk of disease transmission (Murphy and Lugo, 1986; Portillo-Quintero et al., 2015). These global patterns hold true in Hawai‘i; most original tropical dry forests in Hawai‘i have been destroyed (Cordell and Sandquist, 2008) and they are continuously exposed to various threats, mainly due to human activity (Miles et al., 2006). In Hawai‘i, the leading causes for deforestation of drylands include conversion of former dry forests to agricultural-or pastureland, human development, invasion by nonnative species, and wildfires (D’Antonio and Vitousek, 1992).

While ferns are typically found in moist ecosystems, many ferns are epiphytes in these ecosystems, which often exposes them to drier and harsher conditions than terrestrial plants in the same ecosystems (Nitta et al., 2020; Watts and Watkins Jr, 2021). Additionally, some ferns have unique adaptations that allow them to thrive in drylands (Hevly, 1963; Nobel, 1978; Nobel et al., 1978; Sharpe, 2019). In particular, the Cheilanthoideae subfamily of Pteridaceae consists of primarily dry-adapted species worldwide (Gastony and Rollo, 1995; Hevly, 1963). In Hawai‘i, this subfamily is represented by five native species (*Doryopteris* J.Sm. and *Pellaea ternifolia* (Cav.) Link). Taxonomic classification of *Doryopteris* (like much of Cheilanthoideae) has been a challenge for many years because of the lack of clear morphological characters to identify species (see panels B-D in Fig. 1), causing taxonomic uncertainty within the genus (Bouma et al., 2010; Gastony and Rollo, 1995; Ranker et al., 2019a,b; Rothfels et al., 2008; Schuettpelz et al., 2007; Tryon, 1942, 1944; Vernon and Ranker, 2013; Windham et al., 2009; Yesilyurt, 2003; Yesilyurt et al., 2015; Yesilyurt and Schneider, 2010; Zhang et al., 2008, 2007). Despite this uncertainty, to date there are four recognized species of *Doryopteris* that are endemic to the Hawaiian islands: *Doryopteris decora* Brack., *Doryopteris decipiens* (Hook.) J.Sm., *Doryopteris angelica* K.Wood & W.H.Wagner, and *Doryopteris takeuchii* (W.H.Wagner) W.H.Wagner (Vernon and Ranker, 2013; Yesilyurt et al., 2015; Yesilyurt, 2005), along with one named hybrid *Doryopteris subdecipiens* W.H.Wagner (Palmer, 2003). Members of this genus reside in warm and dry shrubland, grassland, and forest habitats (Palmer, 2003), and are known to favor rocky substrates within those habitats (Tryon, 1942). *Pellaea ternifolia* (see panel A in Fig. 1) is found in dry high elevation habitats in Hawai‘i but is also native to Central and South America.

**Figure 1:**
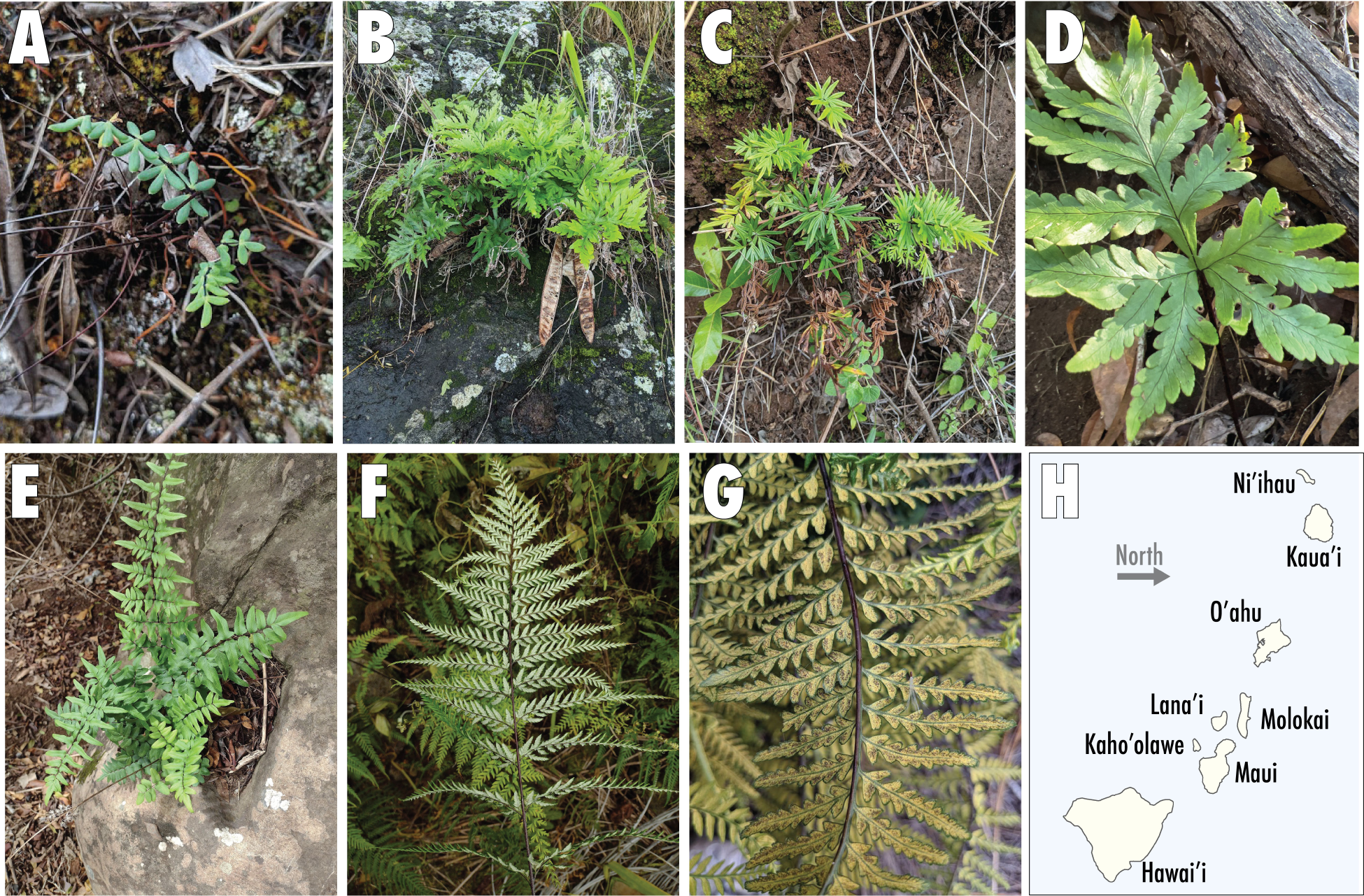
Native (top) and naturalized (bottom) dryland ferns in Hawai‘i. (A) *Pellaea ternifolia*, indigenous to Hawai‘i (Hawai‘i island, Maui, and Kaua‘i) and the Americas, (B) *Doryopteris decipiens*, endemic to Hawai‘i (all major islands), (C) *Doryopteris decora*, endemic to Hawai‘i (all major islands), (D) *Doryopteris angelica*, endemic to Hawai‘i (Kaua‘i single-island endemic), (E) *Cheilanthes viridis*, naturalized, (F) *Pityrogramma calomelanos*, naturalized, (G) *Pityrogramma austroamericana*, naturalized, and (H), map of the major Hawaiian islands. Pictures A, B, C, G by C. M. Tribble, picture D by Susan Fawcett, and pictures E and F by Kevin Faccenda.

Almost 200 native fern species inhabit the Hawaiian Islands, of which 77% are endemic (Palmer, 2003). Geographic ranges of ferns are considered to be strongly determined by habitat availability (Smith, 1993; Tryon, 1986), which may make native fern species particularly susceptible to habitat loss, human-induced habitat disturbance, and competition for resources against naturalized species. For example, naturalized relatives of the Hawaiian members of Cheilanthoideae may be competing for the same dryland habitat that the native species prefer. *Pityrogramma calomelanos* (L.) Link is native to North and South America and the Caribbean (POWO, 2023). *P. austoamericana* Domin is native to Central and South America (POWO, 2023). Both taxa arrived to Hawai‘i in the early 1900s and are now widespread across the islands (see panels F and G in Fig. 1). The first record in Hawai‘i for *P. calomelanos* is in 1903 by Copeland s.n., specimen deposited at MICH, and the first record of *P. austroamericana* is 1903 by Brodie s.n., specimen at BISH. Both *Pityrogramma* species are also part of the Pteridaceae family and share morphological characteristics typical of the Cheilanthoideae subfamily (such as farina, a waxy substance on the abaxial surface of the fronds thought to protect the fronds from damage when desiccated, Kao et al. 2019), although they belong to the subfamily Pteridoideae (Zhang et al., 2017). *Cheilanthes viridis* (Forssk.) Sw. (Cheilanthoideae, Pteridaceae) is another naturalized dryland fern found in Hawai‘i (see panel E in Fig. 1) whose native distribution includes most of eastern and southern Africa and the Middle East (POWO, 2023). *Cheilanthes viridis* is a more recent introduction than *Pityrogramma* spp. and was first recorded in Hawai‘i in 1928 by Lyon s.n., specimen at BISH. *Pityrogramma* spp. and *C. viridis* all meet the criteria for naturalized status, as defined in Brock and Daehler (2020).

Land use has drastically changed over the years in Hawai‘i. In 1848, The Great Māhele (great land division) was one of the first changes in land usage (Chinen, 2020) and resulted in the partitioning of land in the Hawaiian Islands, changing the feudal system to permit land privatization (Kamakau, 1961; Linnekin, 1983). Following the Great Māhele, Hawai‘i became attractive to foreign investors who sought to claim lands in an effort to further develop agricultural plantations. In 1835, the first sugarcane plantation was established in Koloa, Kaua‘i (Hawaiian Sugar Planters Association, 1992) and, by 1890, foreigners and foreign corporations owned three out of four acres of private lands in Hawai‘i (Takaki, 1984). Problems of managing plantations arose soon after the Great Māhele. Sugar, pineapple, and macadamia plantations—often established in former tropical dry forest environments—caused shortages of water in West Maui (Hawaiian Sugar Planters Association, 1992). Since the early 1900s, agricultural corporations and landholders have diverted natural water resources to plantations in former native dryland habitat, resulting in over 90 billion gallons of water being diverted from natural island streams (John A. Engott, 2007). In more recent years, due to changes in the economy, plantations have largely been abandoned (Zou and Bashkin, 1998), and former plantation land is now primarily dominated by invasive grasses such as guinea grass (*Megathyrsus maximus*, Farrant et al., 2023). These land-use changes are a significant conservation concern because naturalized species often invade native communities after some type of disturbance (Smith, 1985). In particular, abandoned drylands create potential hazards to native dryland ferns as naturalized species invade, changing the biotic conditions (naturalized grasses may suppress native species success) and potentially enhancing competition for limited available suitable habitat (naturalized closely related ferns may share similar habitat preferences D’Antonio and Vitousek, 1992).

To better understand the possible adverse effects of (*i*) naturalized species occurrences in the focal native species’ ranges and (*ii*) land-use changes on the survival of native fern species in the Hawaiian islands, we developed an integrated correlative ecological niche modeling (CENM) pipeline that includes information on land use and naturalized taxa. Our study leverages the detailed climate and biodiversity data available for Hawai‘i, and emphasizes the importance of place-based and highly curated approaches to niche modeling for conservation assessments.

## METHODS

Our integrated CENM pipeline leverages highly curated species occurrence and climate data to project suitable area for native species. We first carefully curated occurrence data for seven fern species from herbaria and open-access community science databases and integrated these presence-only data with fine-resolution (8.1 arcsecond spatial units, or approximately 234 × 250 m) climatic variables of the main Hawaiian Islands and a recent CENM (described below Jiménez and Soberón, 2022) to estimate the fundamental niches of the native ferns. The recently developed CENM approach proposed by Jiménez and Soberón (2022) takes as input georeferenced occurrence data as well as randomly sampled points that represent the all the climatic conditions present in the area the focal taxon has access to (the study area, in our case the main Hawaiian islands). The method calculates probabilities of climatic tolerance across that study area and then uses the estimated probabilities to project those probabilities back into geographic space via suitability maps (these steps are explained in the subsequent “METHODS: Climatic niche modeling of native species” section, however see Jiménez and Soberón, 2022, for specific details on the statistical aspects and additional assumptions of the model). We additionally curated data of three naturalized fern species and quantified the degree to which these naturalized taxa occupy the potential ranges of the native ferns. Finally, using a land-use map of the Hawaiian islands, we identified regions with different degrees of disturbance that hold suitable climatic conditions for the native fern species to evaluate the impact of land-use change.

### Species occurrence data

We obtained occurrence records of the native and naturalized species (Fig. 2 step (1)) from digitized museum collections and human observations via the Pteridophyte Collections Consortium (PCC Consortium, 2022), the Global Biodiversity Information Facility (GBIF, 2022), the iNaturalist website https://www.naturalist.org/), the Herbarium Pacificum (BISH) at Bishop Museum, and the Joseph F. Rock Herbarium (HAW) at the University of Hawai‘i at Mānoa. We were unable to recover sufficient accurate occurrences to include *Doryopteris takeuchii* or *Doryopteris subdecipiens* in the study, so all downstream analyses focused on the remaining three *Doryopteris* and *P. ternifolia*. Coordinates for *D. angelica* were obscured on iNaturalist because it is a protected species, so we obtained coordinates for *D. angelica* by reaching out to individual observers. In efforts to obtain more occurrences for *D. angelica*, *D. decora*, *P. calomelanos*, and *P. austroamericana*, we received occurrences from the herbaria BISH and HAW. HAW provided access to digitized collections through the database of the Consortium of Pacific Herbaria, and we requested occurrence records from BISH.

**Figure 2:**
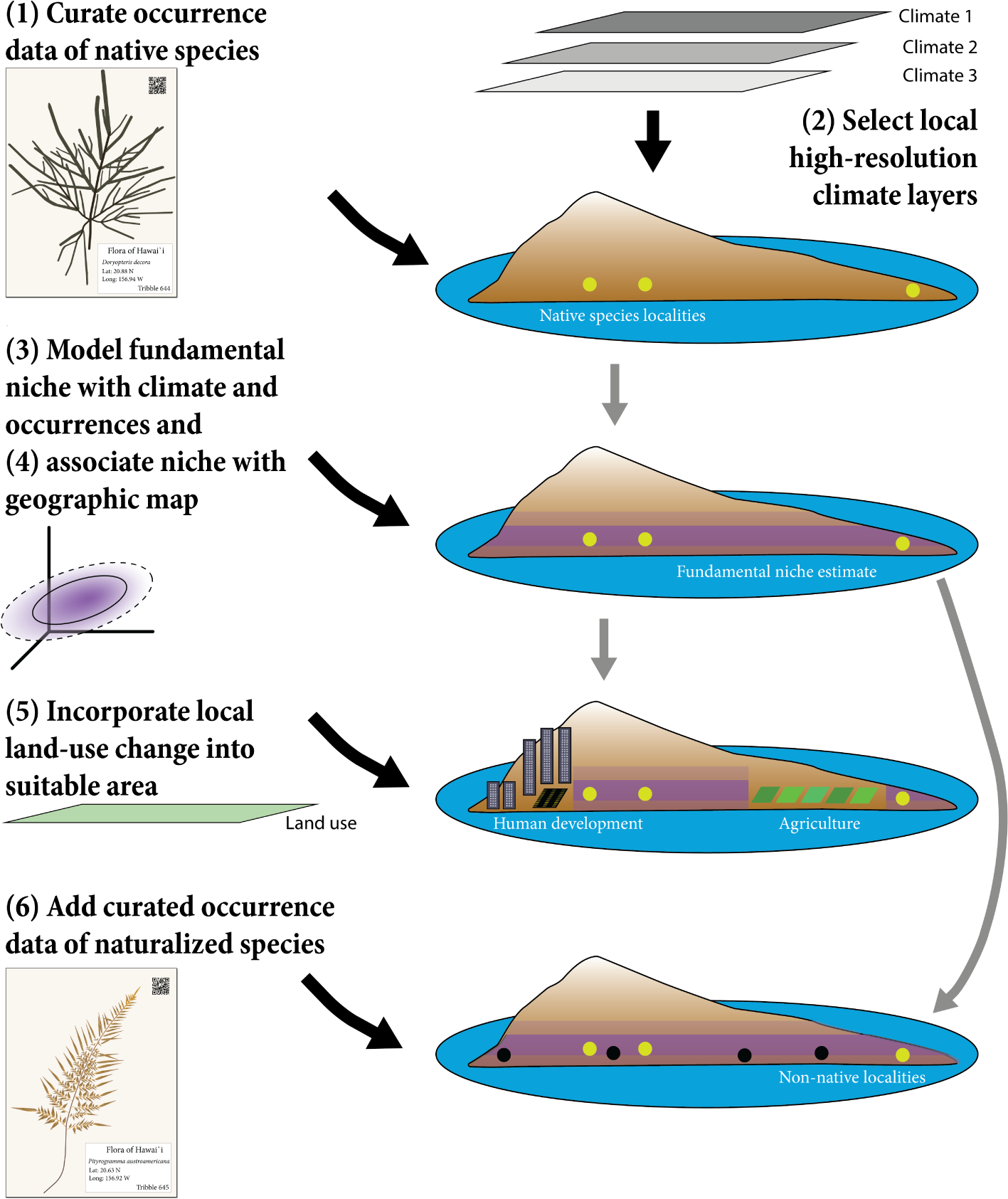
Schematic representation of the analytical workflow. (1) Curate occurrence data of data species. Data from verified physical vouchers is preferred, but iNaturalist or other participatory science databases can supplement if necessary. Curation should include both expert verification of species identifications and careful data cleaning to remove untrustworthy or error-prone GPS data. (2) Gather and transform high-resolution climatic variables, ensuring the variables are not highly correlated and appear sufficient to describe important aspects of the species ecology. (3) Use the data from steps (1) and (2) to estimate the fundamental niche of the native species and (4) identify their potential distributions. The suitability maps created in step (3) were used to: (5) quantify how much suitable area is no longer available to native ferns due to land-use and (6) calculate the percentage of naturalized fern occurrences inside the suitable area.

We then carefully curated the occurrence data through a series of data cleaning steps. First, we separately curated each source’s occurrence records by removing records without GPS coordinates or with high expressed uncertainty in the GPS record (e.g., iNaturalist records with large uncertainty to obscure precise GPS point). We then combined each species’ occurrence records (retaining only pertinent information to the model: species ID, latitude, longitude, source ID, and year) from all sources (PCC, GBIF, herbarium records, and iNaturalist) and removed duplicate records based on matching latitude and longitude. We then removed older occurrences—those recorded prior to 1950—because older records are more susceptible to GPS error and there could be a mismatch between the modern climate data (described below) and the older observations. However, in one case—*D. decora*—we included older observations due to low sample size for this species. Additionally, we included some localities with unknown years to increase our sample size. We plotted the occurrences over a shapefile of the Hawaiian Islands and discarded points located in the ocean or outside of Hawai‘i. To “gut check” the accuracy of our dataset, we mapped the remaining coordinates and closely examined any records that appeared outside of the species’ expected ranges (as defined in Palmer, 2003). Lastly, we examined all physical vouchers or photographs (for iNaturalist records) and confirmed species level identifications by consulting taxonomic experts, including the authors of this study.

Hybridization is hypothesized to occur in *Doryopteris* (especially between *D. decora* and *D. decipiens*, see Palmer, 2003), so we removed several observations with intermediate morphologies. These final taxonomic curation steps are particularly important to accurate niche modeling, as misidentifications can confound estimates of a species’ suitable area (Costa et al., 2015). These cleaning steps resulted in relatively small datasets for each species, so we did not perform additional steps such as spatially thinning the data.

### Climatic data

To model the fundamental niche of the four focal native species in our study, we selected climatic variables that describe factors we believe are likely to influence species ranges (described in Fig. 2 step (2)). We gathered seven environmental variables from the Climate of Hawai‘i website (Giambelluca et al., 2014), which includes detailed breakdowns of climatic and environmental variables: diffuse radiation (W/m²), leaf area index (a proxy for shade/ light availability), relative humidity (%), soil evaporation (mm), solar radiation (W/m²), available soil moisture, actual evapotranspiration (mm). However, the most up-to-date data on Hawaiian temperature and precipitation is available at the Hawai‘i Climate Data Portal (https://www.hawaii.edu/climate-data-portal/). We obtained monthly temperature minimum and maximum values and monthly precipitation values from the Data Portal and transformed those data into five additional variables: mean minimum temperature of the coldest four months (cold mean; *^◦^C*), mean maximum temperature of the hottest four months (hot mean; *C*), mean precipitation of the wettest three months (wet mean; mm), mean precipitation of the driest three months (dry mean; mm), and precipitation seasonality (standard deviation of the monthly precipitation values). All climatic variables were the same resolution (8.1 arcseconds) and projection (WGS84).

The raster layers cover all major Hawaiian Islands except for Ni‘ihau, which was excluded from our analyses. As many of the 12 variables may be highly correlated, we estimated the Pearson correlation coefficient (*r*) for all pairwise comparisons and retained six variables such that all pairwise *r* values satisfied *−*0.7 *> r >* 0.7 (retained variables listed in RESULTS). We decided which variable to drop in order to minimize the overall degree of correlation among variables and to retain variables particularly relevant to the dryland ecology of the focal taxa (e.g., mean precipitation of the driest three months).

### Climatic niche modeling of native species

We applied a methodology that enables the identification of the potential distribution of a species (i.e., the set of sites with suitable climatic conditions where the species can inhabit) by modeling its fundamental niche (*sensu* Soberón, 2007) in a multivariate space defined by climatic variables assumed to be relevant for the survival of the species. In particular, we used the model proposed by Jiménez and Soberón (2022) to estimate the fundamental niches of *Doryopters decipiens*, *D. decora*, *D. angelica*, and *Pellaea ternifolia* with occurrence data and points selected from the study area, which is delimited by making a hypothesis about the dispersal limitations of the species.

We worked under the assumption that the dispersal abilities of these species allow them to reach any of the main islands within the Hawaiian archipelago, while they cannot disperse to other land areas by natural means. In other words, the main Hawaiian islands (Kaua‘i, O‘ahu, Moloka‘i, Lāna‘i, Kaho‘olawe, Maui, and Hawai‘i) represent our hypothesis about the dispersal abilities of the species (known as the movement hypothesis; see Soberón, 2010).

The Jiménez and Soberón (2022) modeling approach assumes that the fundamental niche of the species has an ellipsoidal shape given by a multivariate normal distribution in environmental space which is described through two parameters of interest: a vector that represents both the optimal environmental conditions for the species and the center of the ellipsoid, and a variance-covariance matrix that defines the shape and size of the niche. To estimate the fundamental niche of each native fern species under this approach, we supplied the likelihood function of the model with the curated occurrence data and a sample of 10,000 random points (after extracting their corresponding climatic conditions) within the main Hawaiian islands, representing the set of all available climatic conditions. We then estimated the parameters of interest through a maximum likelihood approach (step 3 in Fig. 2).

The CENM proposed by Jiménez and Soberón (2022) estimates ellipsoids in multidimensional space where the number of dimensions is equal to the number of climatic layers provided to the model and the number of parameters increases at a greater than linear rate. Thus, estimating the parameters of a model with more than three layers is prohibitively computationally demanding. To address this limitation, we created two different models using two sets of three climatic variables each, called Model 1 and Model 2 (variables described in Results).

After obtaining the parameter estimates for each model and species, we created different maps to visualize and compare the outputs of Model 1 and Model 2 as follows (step 4 in Fig. 2). First, we used the estimated parameters to create suitability maps by calculating an index, for each cell, whose values range from zero to one. A suitability index close to 1 means that the environmental conditions associated to that cell are close to the center of the estimated fundamental niche, while a suitability index close to 0 means that the environmental conditions at that cell are either close to the border of the niche or outside of the niche. In each case, we defined the border of the estimated fundamental niche through the 95% confidence ellipsoid of the underlying multivariate normal model. Then, we transformed the suitability maps into binary maps where we assigned a positive value to cells that fall inside the 95% confidence ellipsoid, and a value of zero to cells outside the niche (whose suitability indices are close to zero). For each species, we combined the binary maps for Model 1 and Model 2 by adding the corresponding raster files. With this, we obtained maps whose cells were grouped into four categories: (*i*) *unsuitable*, these correspond to sites with environmental conditions that are outside of the estimated fundamental niches under both Model 1 and Model 2; (*ii*) *suitable under Model 1*, containing sites inside the fundamental niche estimated under Model 1; (*iii*) *suitable under Model 2*, sites inside the fundamental niche estimated under Model 2; and (*vi*) *suitable under Models 1-2*, representing sites predicted as part of the fundamental niche under both Model 1 and Model 2. For further analyses, we only kept regions that were part of the last category.

### Impact of naturalized species on projected species ranges

We created new maps to evaluate to what extent the naturalized ferns, *Pityrogramma calomelanos*, *P. austroamericana*, and *Cheilanthes viridis*, occupy regions identified as the suitable area of the native species. For each native species, we selected the regions identified as suitable under both Model 1 and Model 2 in the previous part of the analyses. We plotted those regions into a new map and added the occurrence data of the naturalized ferns on top (step 6 in Fig. 2).

To quantify the extent to which the native species suitable areas overlap with naturalized species occurrences, we calculated the percentage—for each native species—of all naturalized species occurrences (190 total) that fall within the native species’ suitable area. This metric does not examine overlap between native and naturalized occurrences, but rather between the native’s suitable area (projected suitability based on occurrences) and occurrences of naturalized taxa; a higher percentage of this metric indicates more overlap. We also generated a null expectation for that overlap based on random sampling. For each native species, we randomly sampled 190 points in the study area (main Hawaiian islands, excluding Ni‘ihau). We calculated the percentage of those points that overlapped with the native species’ suitable area (M1 & M2). We repeated this procedure 1000 times to generate a distribution of our null expectation for overlap. In other words, the distribution represents the probability that naturalized species occurrences will overlap with the native species suitable area by chance alone, given the spatial distribution of the native species in the Hawaiian Islands.

### Impact of land-use change on projected species ranges

We downloaded a raster file with a land-cover map provided by the U.S. Geological Survey (Jacobi et al., 2017), which was developed as part of a comprehensive assessment of carbon sequestration potential by natural ecosystems in the State of Hawaii (Selmants et al., 2017). This raster layer depicts the land cover and degree of human disturbance to plant communities on the seven main Hawaiian Islands and is the same resolution (8.1 arcseconds) and projection (WGS84) as the climatic data. Each cell in the raster was classified in a hierarchical fashion within 48 detailed land-cover units, which were also grouped into 27 general land-cover units, 13 biome units, and 7 major land-cover units. We reclassified the raster and created two new land-cover classifications (see Table S3 in Supplementary Appendix), one where we recombined the biome and general land-cover units into 18 new classes. For the second classification, we considered four disturbance regimes (from least to most disturbed): native-dominant, mixed (native-naturalized), naturalized-dominant, and developed.

We calculated the intersection between each of the new land-cover classifications and the binary maps indicating which cells were predicted as part of the fundamental niche of each species under Models 1 and 2 (step 5 in Fig. 2). For this, we first projected and cropped the land-cover rasters so the extent and resolution match the binary maps. We then combined the land-cover and binary rasters into a single raster layer. With the resulting raster, we calculated the proportion of area (relative to the total area predicted as suitable by Models 1-2) covered by the intersection of the suitable areas and each of the land-cover classes. In this way, we were able to quantify how much land in the main Hawaiian islands is climatically suitable for each native fern but is no longer available due land-use changes (Fig. 5).

### Code availability

All the analyses described above were performed using R code (R Core Team, 2020), which we share through this GitHub repository: https://github.com/LauraJim/Hawaiian-Ferns. More information regarding the packages and functions we used is provided in the GitHub repository and the Supplementary Appendix.

## RESULTS

### Data cleaning

We obtained a total of 25,244 occurrences for the seven (four native and three naturalized) species of interest from all the browsed databases. After data cleaning and filtering, we retained 342 occurrences (shown in Fig. 4). This final dataset includes 55 occurrences of *P. calomelanos*, 82 of *P. austroamericana*, 53 of *C. viridis*, 79 of *P. ternifolia*, 29 of *D. angelica*, 45 of *D. decipiens*, and 15 of *D. decora*. These data include occurrences amalgamated by GBIF (2022a,b,c,d,e,f,g,h).

We retained six out of 12 original variables (after removing correlated variables): diffuse radiation (W/m²), leaf area index, relative humidity (%), maximum mean temperature of the hottest four months (called hot mean; *^◦^C*), mean precipitation of the driest three months (called dry mean; mm), and precipitation seasonality (standard deviation of the monthly precipitation values). Figure S1 shows the pairwise correlations between the original 12 variables and Figure S2 shows the pairwise correlations for the selected subset of climatic variables. We sorted these six variables into two models. In Model 1, the niche axes are relative humidity, temperature seasonality, and diffuse radiation. In Model 2, the axes are hot mean, leaf area index, and the dry mean.

### Fundamental niche of native species

Parameter estimates for the two estimated models (Model 1 and Model 2) are provided in Table S1 in the Supplementary Appendix. The estimated ellipsoids of *D. angelica* were the smallest and, under Model 1, its fundamental niche is nested inside ellipsoids representing the fundamental niches of *D. decipiens* and *D. decora* (see Fig. S5 in the Supplementary Appendix). Under both models, the estimated niches of the ferns in *Doryopteris* are similar to each other—their centers are close and they intersect (see Figures S5 and S6). For *P. ternifolia*, the estimated optimal conditions (ellipsoid’s center) were very different from the ones estimated for *Doryopteris* spp. under Model 2. This species’ fundamental niche includes colder temperatures and higher leaf area indices than other taxa. However, under Model 1, *P. ternifolia*’s fundamental niche is also quite broad and contains the ellipsoids that represent the niches of the other native species. In the Supplementary Appendix, we also provide the suitability maps, for each species and model, that were calculated with the parameter estimates (see Fig. S7).

After identifying the 95% confidence ellipsoids of the underlying model for the fundamental niche, we obtained binary maps for Model 1 and Model 2, separately. Each pair of binary maps was then combined to obtain the maps shown in Figure 3. All the main Hawaiian islands hold climatically suitable area for the four native species (except Kaho‘olawe with *D. decora*). For *D. angelica*, only Kaua‘i contains regions identified as suitable under both Model 1 and Model 2.

**Figure 3:**
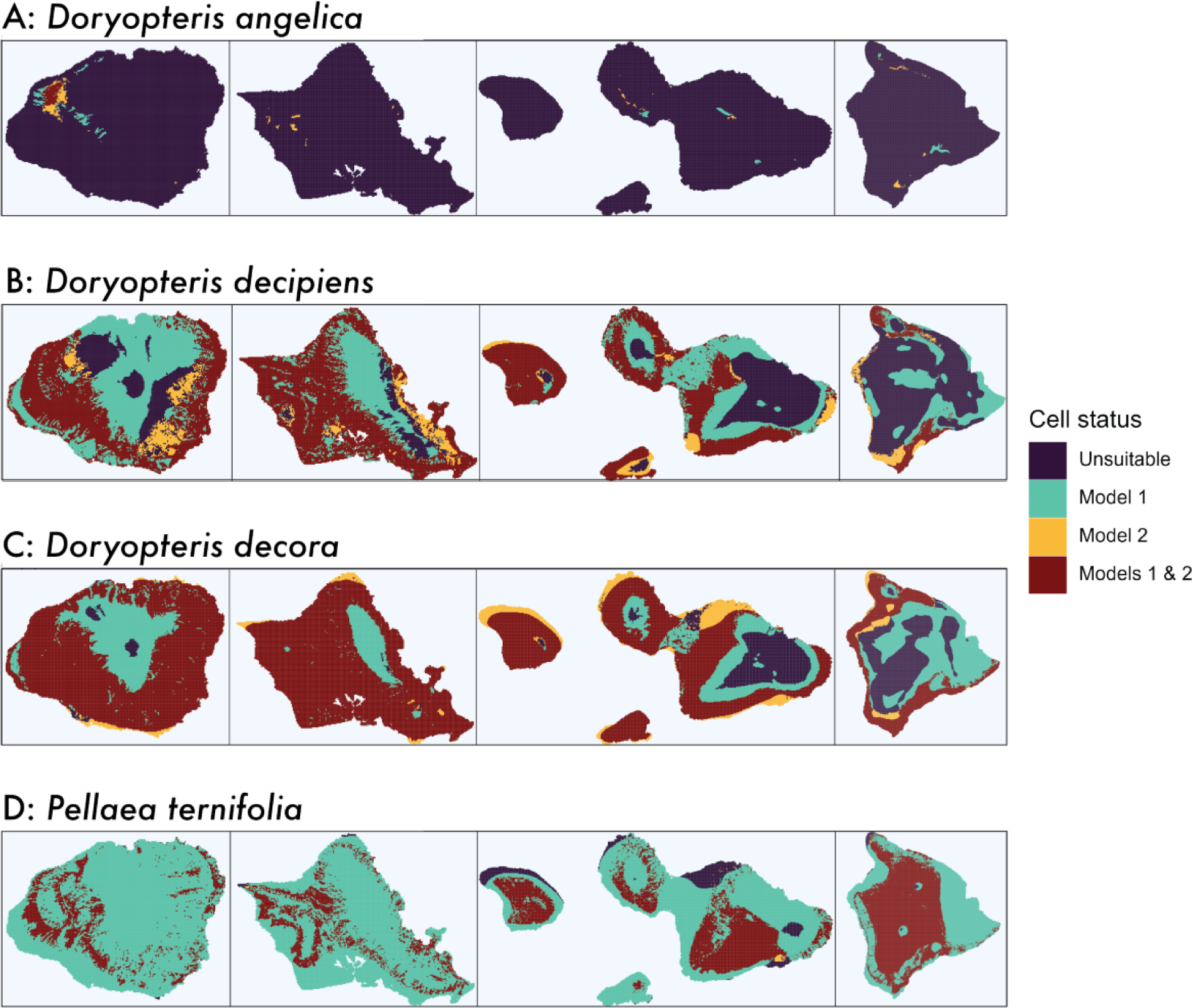
Suitable areas under Model 1 (teal), Model 2 (yellow), and the overlap of 1 and 2 (burgundy) for native Hawaiian dryland ferns: (A) *Doryopteris angelica*, (B) *Doryopteris decipiens*, (C) *Doryopteris decora*, and (D) *Pellaea ternifolia*.

Under Model 1, *P. ternifolia* has the largest potential range, however, after intersecting it with the predicted potential range under Model 2, the final potential range was drastically reduced in Kaua‘i and O‘ahu.

### Impact of naturalized species on projected species ranges

We mapped the suitable area of native species (visualized in Fig. 4 and added the occurrence points of both native (green points) and naturalized ferns on top of these maps (yellow points) to evaluate two questions: (1) Does the suitable area include sites where the species was already identified as present? and (2) To what degree do the naturalized ferns occupy the potential distribution of the native ferns? *D. angelica* has the smallest suitable area of native species within the main Hawaiian islands, where only Kaua‘i (left panel in Fig. 4) holds meaningful sites with suitable conditions for this species. The suitable areas of *D. decipiens*, *D. decora*, and *P. ternifolia* are wider and all the main islands have regions with suitable conditions (except Kaho‘olawe with *D. decora*).

**Figure 4:**
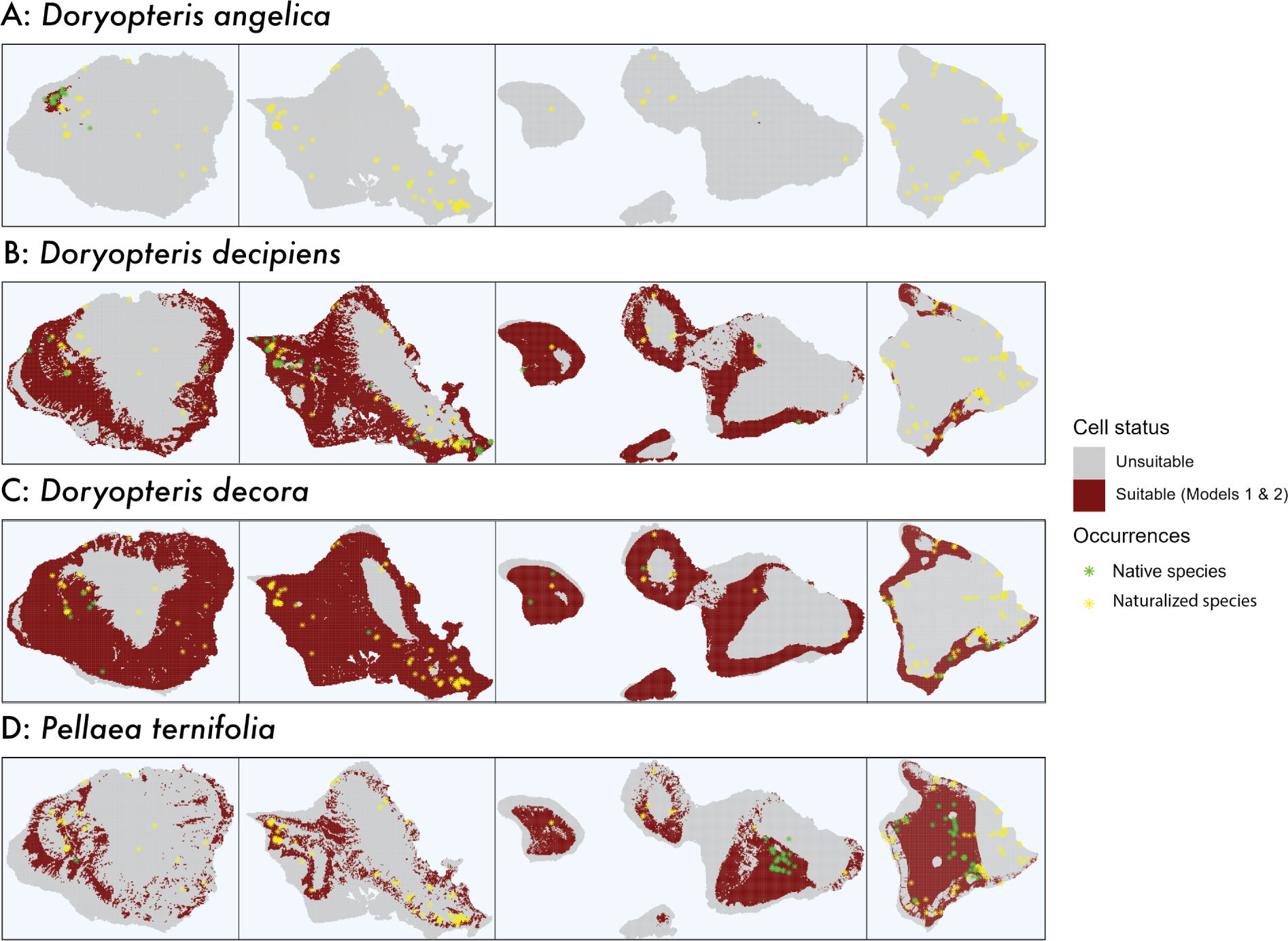
The binary maps obtained for each native species. The regions colored in burgundy represent cells that were identified as part of the fundamental niche of the species under both Model 1 and Model 2, which we refer to as the suitable area for the species. Panels show suitable area for:(A) *Doryopteris angelica*, (B) *Doryopteris decipiens*, (C) *Doryopteris decora*, and (D) *Pellaea ternifolia*. Green asterisks represent occurrences of the respective native species, while yellow asterisks represent occurrences of all studied naturalized relatives.

The suitable area of *D. angelica* correctly predicted 82.75% of the conspecific occurrence points, while 2.6% of the naturalized ferns occurrence points are inside this range (Table 1). For *D. decipiens*, the suitable area included 91.1% of the conspecific native occurrences and 42.3% of the naturalized occurrences are inside the species’ suitable area (Table 1). 100% of the occurrences identified as *D. decora* are inside its corresponding suitable area, which contains the highest percentage of naturalized occurrences at 65.6% (Table 1). Finally, *P. ternifolia*’s suitable area contained 96.2% of the occurrences and 34.3% of the naturalized occurrences (Table 1).

**Table 1:**
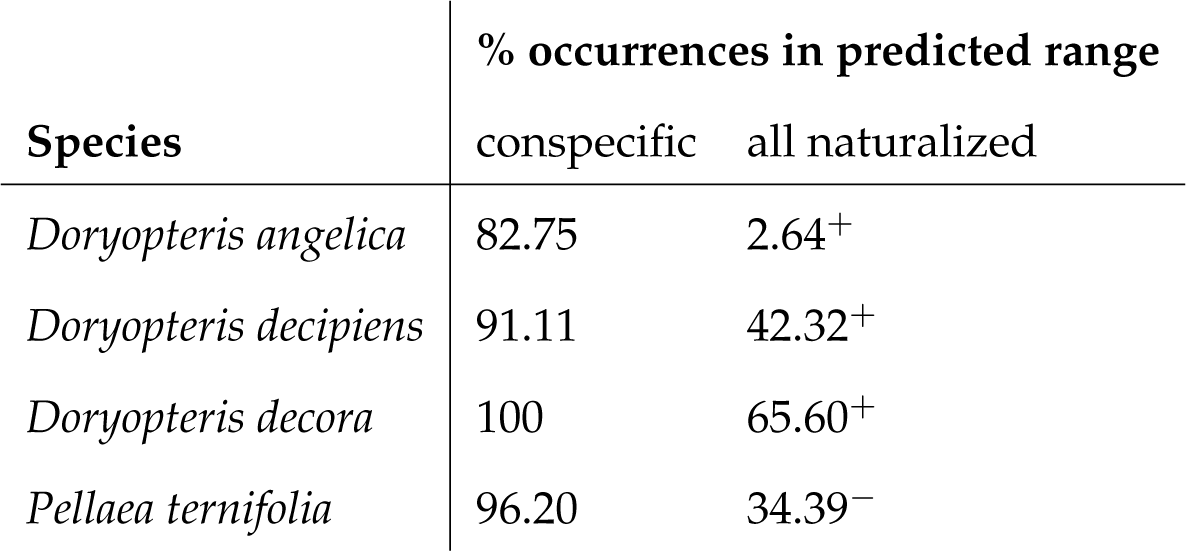
Percent of occurrences of the listed conspecific native species (e.g., *Doryopteris decora* for row *Doryopteris decora*) and for all naturalized species that fall within the predicted ranges for native species. A higher value for conspecific occurrences indicates that the predicted range includes more of that species’ occurrences, whereas a higher value for all naturalized occurrences indicates that the predicted range includes more occurrences of naturalized species. Superscript symbols (+ vs. -) indicate whether the percent overlap of naturalized occurrences in the native range is greater or lower—respectively—than expected by chance (see also Fig. S9 in the Supplementary Appendix).

| Species | % occurrences in predicted range |  |
| --- | --- | --- |
|  | conspecific | all naturalized |
| *Doryopteris angelica* | 82.75 | 2.64<sup>+</sup> |
| *Doryopteris decipiens* | 91.11 | 42.32<sup>+</sup> |
| *Doryopteris decora* | 100 | 65.60<sup>+</sup> |
| *Pellaea ternifolia* | 96.20 | 34.39<sup>-</sup> |

For all native species, the percent overlap with naturalized occurrences are significantly different than the null expectation generated through random sampling (Fig. S9). All *Doryopteris* species have higher overlap than expected by chance (Fig. S9A–C), suggesting that these species frequently co-occur with the naturalized ferns in this study. However *P. ternifolia* has lower overlap than expected by chance (Fig. S9D), suggesting that this native species occupies significantly different areas than the studied naturalized taxa.

### Impact of land-use change on projected species ranges

We overlaid the potential distribution (under Model 1 and Model 2) of each species with the land-cover layer to partition the climatically suitable areas into land-use types. Under the land-use classification based on biomes and general land-use units, 17 out of the 18 categories intersected with the suitable area of all species (coastal strand is the only land-cover type the does not intersect with any species’ suitable area; Fig. 5 and Table S2 in the Supplementary Appendix). For *D. angelica*, 80.6% of the suitable area is covered by mesic forest. About 29% of the potential distribution of *D. decipiens* presents some degree of development or is not vegetated. For both *D. decipiens* and *D. decora*, about 6% of their potential distribution is currently used for agriculture. About a quarter of *P. ternifolia*’ potential distribution is not vegetated (likely because of this species’ affinity for high elevation volcanic habitat with limited plant cover).

**Figure 5:**
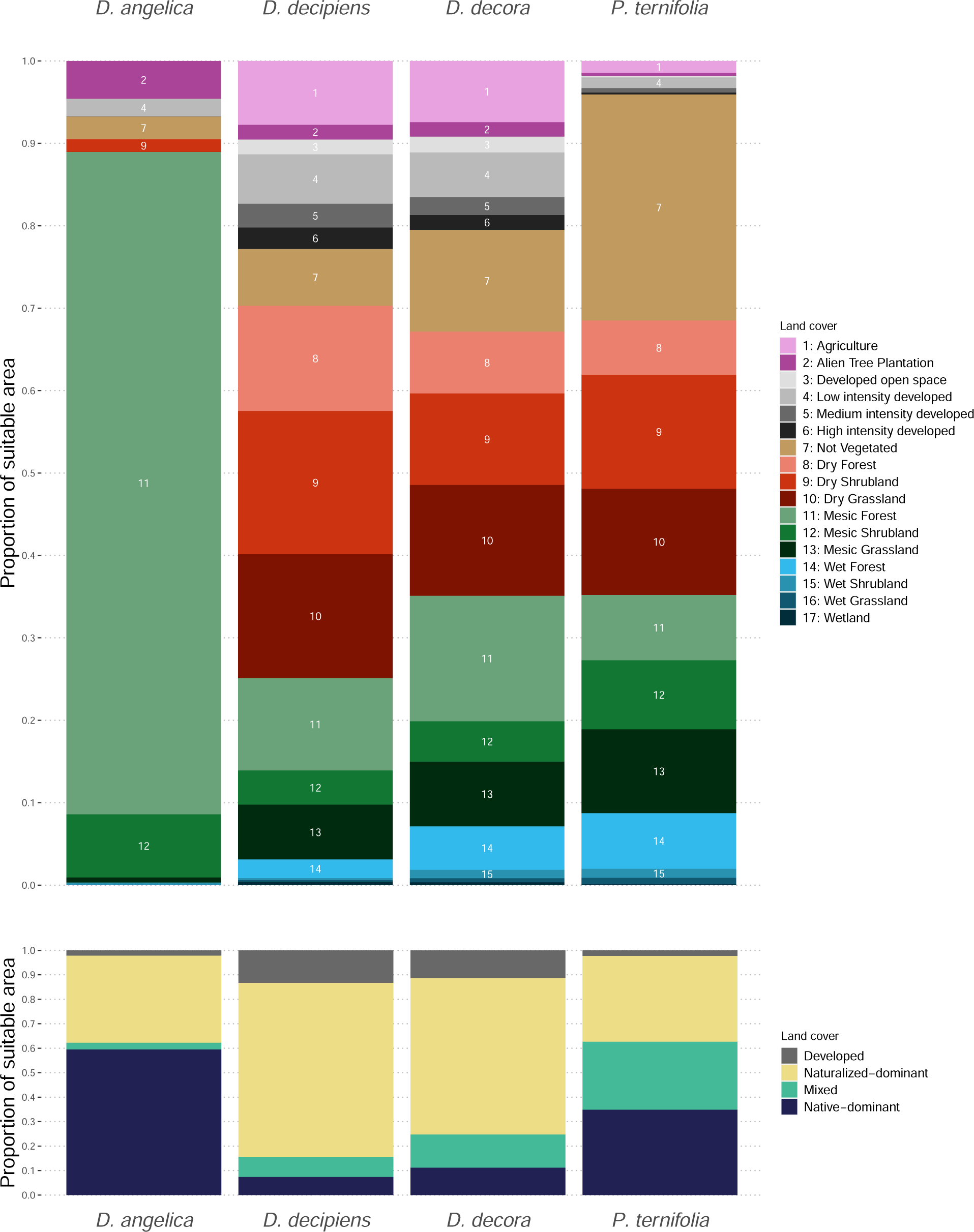
Land uses of inferred climatically suitable areas for native species species. The barplots show the proportion of the total suitable area (sites inside the 95% ellipsoids representing the fundamental niche) that is covered by each land-use type in the corresponding classification. The first classification combines biomes and general land-cover units and the second classification includes four disturbance regimes from the most disturbed (Developed) to the most conserved

Under the second classification, *D. angelica*’s potential distribution is mainly covered by native-dominant (58.5%) and naturalized-dominant (35.2%) habitats (see Fig. 5 and Table S2 in the Supplementary Appendix). Developed (14%) and naturalized-dominant (69.3%) habitats cover more than 80% of *D. decipiens*’s potential distribution, while only 6.9% of the suitable area is represented by native habitats. We observed a similar pattern for *D. decora* where only a small fraction of the suitable area is covered by native-dominant habitats (16.9%). However, *P. ternifolia*’s potential range still presents a native habitat coverage of 37.8%, while the remaining area is mainly covered by mixed (27.9%) and naturalized-dominant habitats (32.4%).

## DISCUSSION

Hawai‘i is a longstanding model for understanding ecological and evolutionary processes, and it is exceedingly important to continue studying rapid changes on the islands in the face of the global biodiversity crisis. The rate of naturalization in Hawai‘i is increasing, and nonnative plants now exceed natives in the Hawaiian flora (Brock and Daehler, 2021). Naturalized plant species may have a number of different impacts on native ecological processes. They can compete with native species for water or nutrients and act as primary or alternate hosts for pests and diseases (Chau et al. 2013; Garcia-Serrano et al. 2007; Mason et al. 2012, but see Westerband et al. 2021), and fern gametophytes may have interspecific competitive effects (Testo and Watkins Jr, 2013). However, a common impact is physical displacement when naturalized species colonize disturbed sites and occupy them before the native species can reestablish. Similarly, land-use changes such as agriculture and human development may render climatically appropriate habitat unavailable to native species. Our pipeline for modeling species fundamental niche while addressing the potential impacts of naturalized species and land-use change demonstrates the importance to conservation efforts of modeling multiple dynamic threats to biodiversity, especially in the highly disturbed habitat of Hawai‘i dryland ecosystems. We illustrate this pipeline using ferns—plants that tend to not be dispersal-limited given the ability of wind-dispersed spores to travel long distances (Schaefer and Fontaneto, 2011) and thus whose distributions are more likely to be shaped by abiotic and/or biotic factors of their environments, rather than by dispersal limitation.

### Importance of land-use data for predicting species’ suitable areas

Our results demonstrate an important lesson for plant conservation: an accurate picture of plants’ conservation statuses must account for present and historical land use. We demonstrate that while native species *Doryopteris decipiens* and *D. decora* have broader fundamental niches—and thus more suitable land area in Hawai‘i—than the congeneric *D. angelica*, much of the land predicted as suitable for *D. decipiens* and *D. decora* is dominated by human development or naturalized plant habitat (Fig. 5). Of the four native species in our study, only one species (*Doryopteris angelica*) had more than half of its suitable area characterized as primarily native habitat, and this species also has by far the smallest suitable area of all natives. Agriculture accounts for a small percentage of the species’ suitable area, though *D. decora* is more affected than the other species. *D. angelica* is considered endangered by the US Fish and Wildlife Service and is the target of several conservation efforts including as Genetic Safety Net Species through the State of Hawai‘i Department of Land and Natural Resources and the Hawai‘i State Wildlife Action Plan (DLNR, 2023). In contrast, conspecifics *D. decora* and *D. decipiens* have much larger suitable areas which are much more dominated by naturalized habitat (native = 6.95% for *D. decora* and native = 16.92% for *D. decipiens*, see Table 1). While these species have much larger potential ranges, the quality of this habitat is potentially lower, presenting additional conservation concerns. Our results thus suggest that—by accounting for land use—additional conservation concern may be warranted for these species. Our pipeline thus provides an avenue for conservationists to quantitatively assess the impact of land use on species of interests, as suggested in Gillson et al. (2020).

We contextualize the temporally static land use change data from (Jacobi et al., 2017) with a deeper understanding of the history of land use change in Hawai‘i. The data generated by Jacobi et al. (2017) represent a snapshot of Hawai‘i in 2014 when former agricultural lands had been mostly abandoned and had converted—largely through neglect—to naturalized grass-dominated landscapes. The low proportion of agricultural land projected as suitable for the species of our study is likely an artifact of this history. This history also illuminates possible conservation futures, and we echo calls for fallowed agricultural land to be targeted for Hawaiian dry forest conservation, including of the native fern flora; restoration of fallow agricultural land has also been suggested as a fire recovery and prevention mechanism (Bond-Smith et al., 2023; Trauernicht et al., 2018).

### Impact of naturalized ferns

*Doryopteris decora* is the native fern whose predicted range includes the highest proportion of naturalized occurrences (65.6% of all naturalized occurrences in Hawai‘i overlap with the predicted range, Table 1), suggested that this species is the most potentially impacted by competition from related naturalized ferns. We emphasize that our study does not demonstrate that naturalized ferns are directly competing with natives, nor that these naturalized taxa have negative impacts on native species. Rather, we show that naturalized ferns indeed share very similar climatic preferences to native ferns and occur in habitats predicted to be suitable for natives, especially of *D. decora*. While it is possible that *D. decora* persists in these environments despite co-occurrence with naturalized taxa, our difficulty in obtaining recent and accurate occurrence data for this species (see METHODS: Species occurrence data) suggests that *D. decora* may be failing to thrive in an invaded, altered landscape. We suggest that further conservation attention be given to this species, and follow-up studies should test for evidence of competition between *D. decora* especially and naturalized dryland ferns through controlled greenhouse experiments or field trials that evaluate growth and reproductive success in mixed (native + naturalized) vs. isolated population. While only 2.6% of naturalized occurrences occur in the suitable area for federally listed *D. angelica*, the predicted area itself is significantly smaller than for all other native species in this study. Visual inspect of the occurrences (Fig. 4A) shows many naturalized individuals (yellow) in and around the small suitable area (burgundy) and *D. angelica* occurrences (green), and we demonstrate the 2.6% is significantly higher than our null expectation for overlap (Fig.S9). Competition from naturalized ferns may thus still be a concern for this species.

### Accuracy of modeling approach

For most native species, our modeling approach results in suitable areas that include *>* 90% of observed occurrences. *D. angelica* is the one exception, with only 82.7% of occurrences falling into the suitable area. This may be because *D. angelica* is predicted to have far less suitable area than the other native species, implying that our model is inferring a very specific set of climatic conditions necessary for *D. angelica* to survive. *D. angelica* is known from only a few populations that are clustered in similar regions of Kaua‘i island, and it is possible our model is inferring more strict climatic conditions because the existing individuals and populations are highly clustered in climatic and geographic space.

On the other hand, our suitable area estimate for *Doryopteris decora* (and for *Pellaea ternifolia* under Model 1) is quite broad (Fig. 3). Regions in the suitable area for *D. decora* but where we have no occurrences (e.g., much of O‘ahu) may represent sites where naturalized or native congeneric taxa have competitively excluded the species or where other biotic factors limit the species. For *P. ternifolia*, our results may suggest that Model 1 variables hold little information about suitability: relative humidity, temperature seasonality, and diffuse radiation. In order to maximize comparability across species, we used the same set of climatic variables for all estimates, but future studies on *P. ternifolia* could consider other variables that could be more appropriate for modeling suitability.

The number of occurrences per species used for niche modeling varied from 15 (*D. decora*) to 79 (*P. ternifolia*). It is possible that these small samples sizes may have impacted the accuracy of the resulting niche models (Jiménez-Valverde et al., 2009; Sillero et al., 2021; Wisz et al., 2008), though all taxa but *D. decora* are over the general threshold of 20-30 points (threshold from Sillero et al., 2021), a possible explanation for the breadth of the inferred suitable area for this species.

There exists a trade-off between data curation (via removing questionable observations) and study sample size. We have tended towards data curation in this study given the taxonomic challenges (for *Doryopteris* spp. in particular) and the small spatial scale. However, we have chosen to not spatially thin our data or directly account for sampling biases to not further shrink the sample sizes in this study. While we believe sampling bias is less likely to be a factor in this study than others (given the small spatial scale of the study area, the density of trails or other access across the landscape throughout much of Hawai‘i, among other factors), our occurrences probably fall along trails more often than expected by chance in wilderness areas. If the relatively proportion of trails to land area differs based on environment, it is possible that this bias could effect our estimates of both native suitable area or the overlap of naturalized occurrences with native suitable area.

Our niche modeling approach in this paper uses occurrences and observations of fern sporophytes to infer the species’ suitable areas. However, ferns have a dual life cycle in which both gametophyte and sporophyte exist independently (not nutritionally dependent on each other Krieg and Chambers, 2022; Pinson et al., 2017). For most ferns, the sporophyte is seen as dominant and the gametophyte is reduced in both size and longevity. However, some gametophytes persist for long time periods, occasionally with ranges that differ (or exceed) the range of the sporophyte (Pinson et al., 2017, 2022, though most ferns with long-lived gametophytes are epiphytic and the focal taxa in this study are terrestrial), likely due to differences in the underlying physiological tolerances of gametophytes vs. sporophytes (Krieg and Chambers, 2022). Even when gametophytes are short lived, they may differ from sporophytes in key physiological limits that impose unexpected bounds on the responses of fern species to land-used change or naturalized species competition. Thus, follow-up studies should investigate any differences in physiological tolerances of gametophyte vs. sporophyte dryland ferns.

### Applicability of pipeline to future studies

The pipeline we outline in this study (illustrated in Fig. 2) may guide future studies that use CENMs for conservation applications. Here, we outline three main takeaways for researchers to use moving forward: First, our integration of land use into evaluations of suitable area for natives (step (5) in Fig. 2) requires a place-based understanding of the history behind current land use. We position our study within the context of land-use change in Hawai‘i; land privatization during Great Māhele encouraged the development of plantations in former Hawaiian drylands. Despite the current abandonment of most plantations, much of that land is still unavailable for native species because naturalized species now occupy former native habitat. Our land use change results demonstrate the impact of this history on native dryland plants; only one native species is predicted to have more than 50% climatically suitable area covered by native ecosystems.

Second our approach allows for and promotes the careful curation of both climatic and occurrence data (steps (1) and (2) in Fig. 2). Rather than use a broad set of generic climatic variables (e.g., from WorldClim), we obtained regionally specific and high-resolution climatic layers that describe features likely to be relevant to the species of interest based on their ecology. We encourage researchers to seek out similar datasets whenever possible, though we also acknowledge that many regions of the world do not have access to such data (e.g., see discussion in Soria-Auza et al. 2010).

Third, we emphasize the importance of using a mathematical model that aligns with our theoretical definitions of the niche. Our approach uses the Jiménez and Soberón (2022) model to estimate the fundamental niche—the set of climatic conditions that an organism could inhabit in the absence of biotic interactions. Thus we must also evaluate the potential role of other species. In our case we set out to evaluate how naturalized taxa may impose additional restrictions on where natives occur. We find evidence suggesting that naturalized species often occur in the suitable area for native species (particularly with *D. decora*), consistent with competition for limited available dryland habitat. We encourage researchers to ensure that their model of choice aligns with their theoretical understanding of the ecological niche.

### Conclusions

Our novel pipeline unites new developments in correlative ecological niche modeling with a place-based understanding of land-use change and potential competition from naturalized species. We demonstrate that native Hawaiian dryland ferns may be competing for limited suitable habitat with naturalized relatives, and our results point to the drastic reduction in available area due to land-use changes such as human development and non-native ecosystems, especially fallowed agricultural lands. We advocate for continued conservation efforts for the endangered *Doryopteris angelica*, additional conservation assessment for *D. decora*, and restoration of fallowed agricultural lands to build up more native habitat. Our study also demonstrates the value of our integrative approach and serves as a model for multidimensional conservation assessments based on correlative ecological niche modeling.

## Acknowledgements

K.E.C. received support from the DNA-Based Discoveries in Hawai‘i’s Biodiversity Research Experience for Undergraduates (REU) Program at the University of Hawai‘i at Mānoa (NSF DBI #1950950) and from NSF DBI # 2135175 awarded to K.H. L.J. received partial support from Agencia Nacional de Investigación y Desarrollo (ANID-Chile) in two modalities: (1) FONDECYT posdoctorado 2023, Folio 3230511, and (2) Centro de Modelamiento Matemático (CMM), BASAL fund FB210005 for centers of excellence. This material is based upon work supported by the NSF Postdoctoral Research Fellowships in Biology Program under Grant No. 2109835 to C.M.T. HAW and BISH directly contributed herbarium specimen data for occurrence records in this study. Alicia Rozet and Christian Tabor graciously provided feedback and assistance with the Hawaiian language abstract translation.

## Author contributions

K.E.C, C.M.T., and R.Z.F conceived the study. K.E.C, L.J, and C.M.T performed analyses and wrote the paper. R.Z.F., M.K.T., and K.H. provided feedback on analyses and edited the paper.

## Data availability statement

All the R code needed to reproduce all our analyses and figures is available at this GitHub repository: https://github.com/LauraJim/Hawaiian-Ferns. The occurrence data for all the species is provided in the same GitHub repository, except for *Doryopteris angelica* given conservation concerns regarding this native Hawaiian species. The rasters with the climate variables and land-use layers can be found at their original sources using the links provided in the Methods section.

