## Supplemental Materials for "Conservation applications of niche modeling: native and naturalized ferns may compete for limited Hawaiian dryland habitat"

#### Selection of climatic variables for ecological niche models

We considered two types of climatic variables in our analyses, one type consisted on generic environmental variables from the Climate of Hawai'i website (http://climate.geography.hawaii.edu/, Giambelluca et al. (2014)), and the second included climatic variables that we created taking into account the natural history of the species of interest. In the first group, we considered the variables: diffuse radiation (W/m²), leaf area index, relative humidity (%), soil evaporation (mm), solar radiation (W/m²), available soil moisture, and actual evapo-transpiration (mm). Then, we created five variables: mean minimum temperature of the coldest four months (cold mean; °C), mean maximum temperature of the hottest four months (hot mean; °C), mean precipitation of the wettest three months (wet mean; mm), mean precipitation of the driest three months (dry mean; mm), and precipitation seasonality (standard deviation of the monthly precipitation values). These variables were calculated from minimum and maximum monthly temperature values and monthly average precipitation values using the most up-to-date data on Hawaiian temperature and precipitation (available at the Hawai'i Climate Data Portal, https://www.hawaii.edu/climate-data-portal/), and functions in the `raster` package v3.6 (Hijmans et al., 2015; R Core Team, 2020). The code used for this part of the analyses is included in the script called "01_Create_climate_layers.R", available at the GitHub repository: https://github.com/LauraJim/Hawaiian-Ferns.git

Figure S1 shows pairwise scatterplots of the climatic conditions observed at the occurrence (colored dots) and random (black dots) points, for each pair of the twelve original variables considered for the ecological niche modeling, and each species of interest. We observed clear correlation patterns between some variables, which were then corroborated through a correlation analysis. Figure S2 shows a similar matrix of scatterplots where only the six less-correlated climatic variables are included. Relative humidity and hot mean still seem to be correlated, however, they were not used in the same ecological niche model.

We used the geographic coordinates of each occurrence point (only for native ferns) to extract their corresponding climatic values using the twelve selected variables (see the file called "03_Extracting_climatic_values.R" at the GitHub repository). With this, we calculated the correlations between each pair of climatic variables (see the file called "02_Data_cleaning_Correlation.R" at the GitHub repository) and eliminated from the following analyses the highly correlated variables (whose absolute correlation value was higher than 0.7). Figure S3 shows the resulting correlation values. We selected a subset of six climatic variables: diffuse radiation, relative humidity, precipitation seasonality, leaf area index, hot mean, and dry mean. The first three variables were included in the niche model called Model 1, and the second set of three variables was included in the niche model called Model 2. Figure S4 shows the correlation coefficients calculated between each pair of the selected variables.

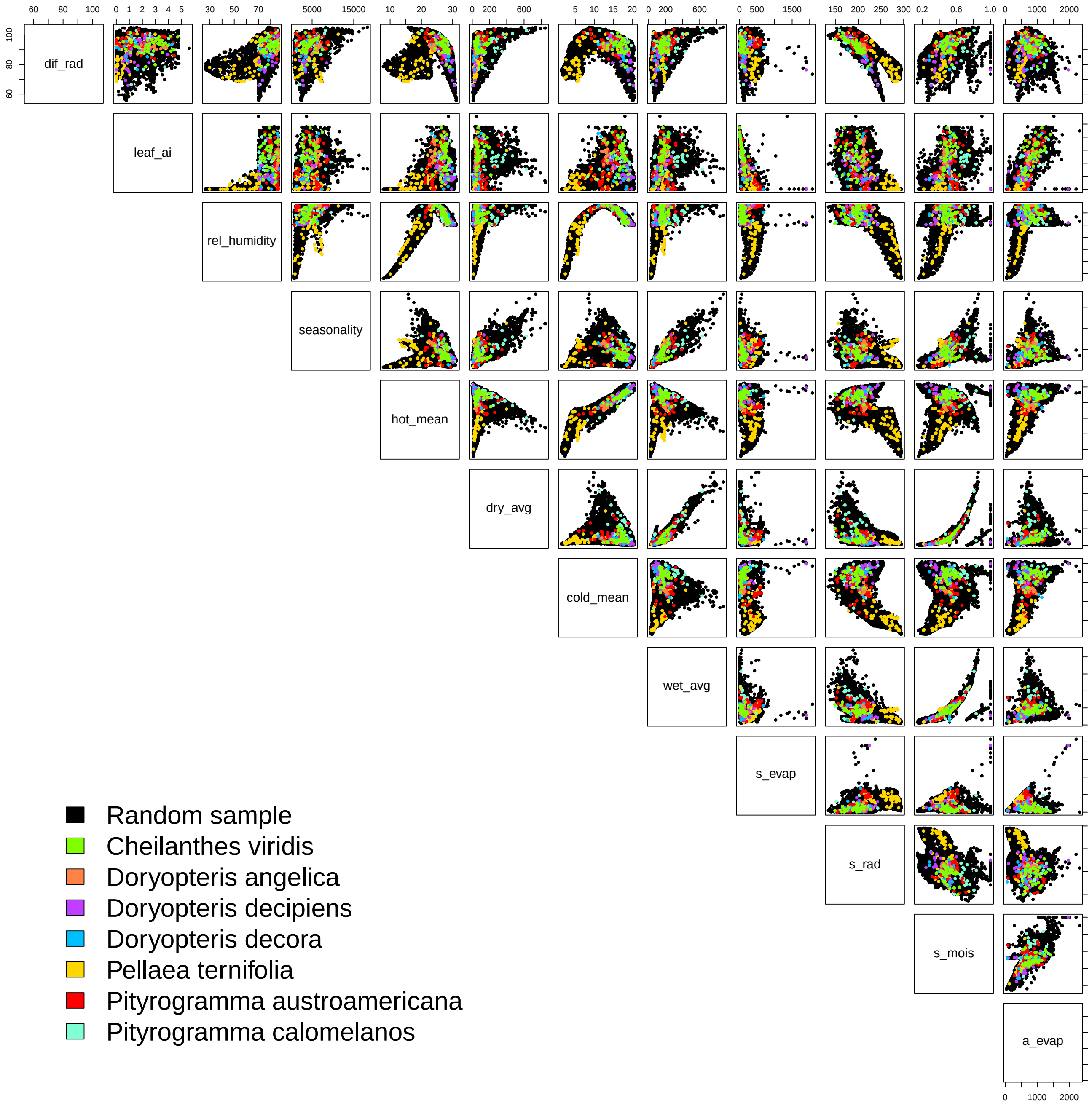


**Figure S1:** Pairwise scatterplots of the original 12 climatic variables. Black points represent 10,000 points randomly sampled from the layers and colored points represent the climate values of individual species occurrences.

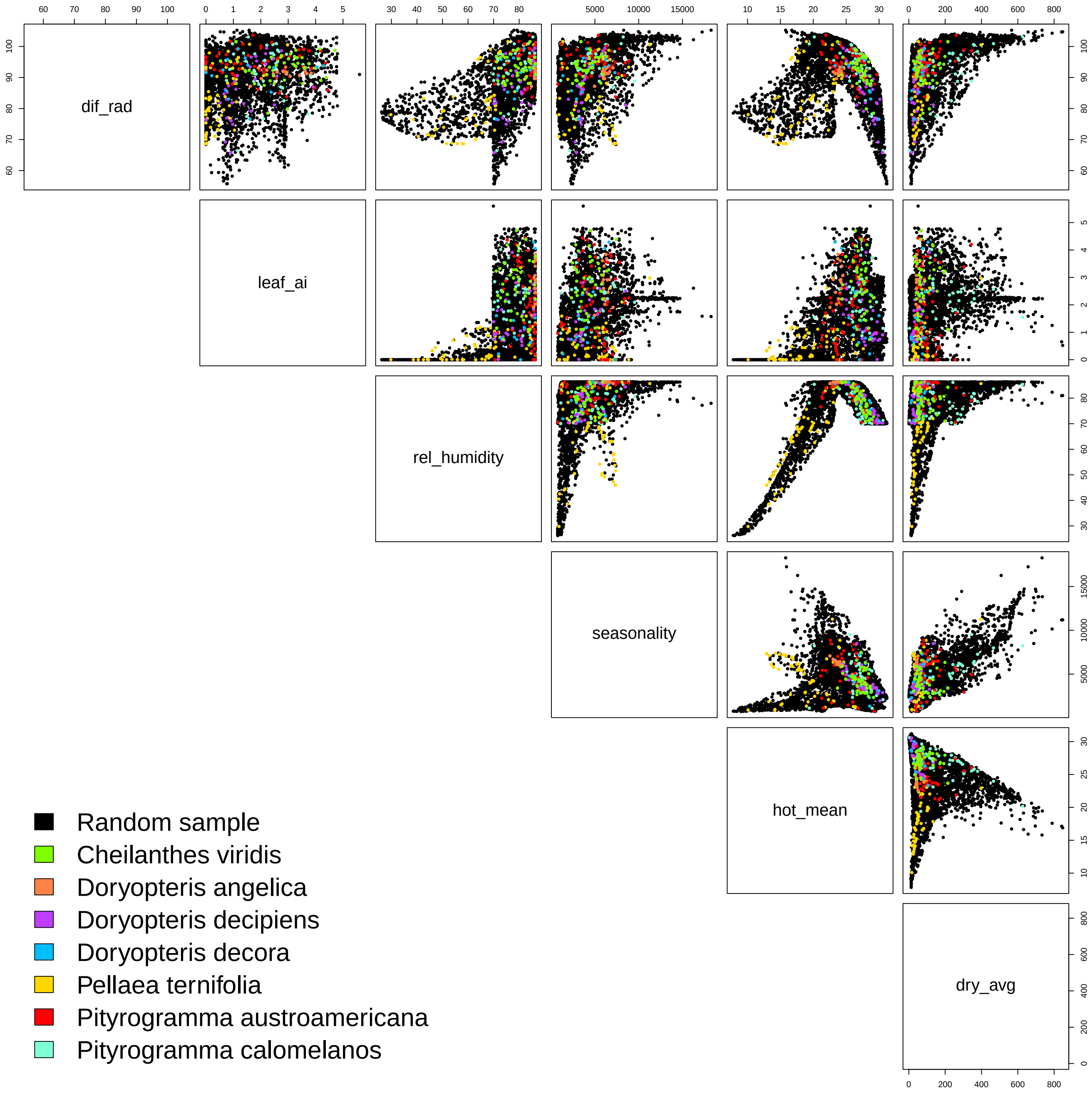


**Figure S2:** Pairwise scatterplots of the selected six climatic variables. Black points represent 10,000 points randomly sampled from the layers and colored points represent the climate values of individual species occurrences.

| | dif_rad | leaf_ai | rel_humidity | seasonality | hot_mean | dry_avg | cold_mean | wet_avg | s_evap | s_rad | s_mois | a_evap |
|---|---|---|---|---|---|---|---|---|---|---|---|---|
| dif_rad | 1.00 | 0.40 | 0.65 | 0.12 | 0.30 | 0.38 | 0.29 | 0.38 | −0.32 | −0.83 | 0.44 | 0.25 |
| leaf_ai | 0.40 | 1.00 | 0.42 | 0.09 | 0.45 | 0.17 | 0.44 | 0.22 | −0.72 | −0.55 | 0.27 | 0.70 |
| rel_humidity | 0.65 | 0.42 | 1.00 | 0.15 | 0.58 | 0.16 | 0.47 | 0.24 | −0.29 | −0.65 | 0.29 | 0.38 |
| seasonality | 0.12 | 0.09 | 0.15 | 1.00 | −0.24 | 0.27 | −0.13 | 0.72 | 0.14 | 0.03 | 0.41 | 0.16 |
| hot_mean | 0.30 | 0.45 | 0.58 | −0.24 | 1.00 | 0.11 | 0.94 | −0.01 | −0.42 | −0.69 | 0.18 | 0.43 |
| dry_avg | 0.38 | 0.17 | 0.16 | 0.27 | 0.11 | 1.00 | 0.14 | 0.82 | −0.09 | −0.37 | 0.79 | 0.26 |
| cold_mean | 0.29 | 0.44 | 0.47 | −0.13 | 0.94 | 0.14 | 1.00 | 0.08 | −0.43 | −0.70 | 0.20 | 0.43 |
| wet_avg | 0.38 | 0.22 | 0.24 | 0.72 | −0.01 | 0.82 | 0.08 | 1.00 | [illegible] | −0.30 | 0.80 | 0.35 |
| s_evap | −0.32 | −0.72 | −0.29 | 0.14 | −0.42 | −0.09 | −0.43 | [illegible] | 1.00 | 0.51 | 0.12 | −0.21 |
| s_rad | −0.83 | −0.55 | −0.65 | 0.03 | −0.69 | −0.37 | −0.70 | −0.30 | 0.51 | 1.00 | −0.40 | −0.36 |
| s_mois | 0.44 | 0.27 | 0.29 | 0.41 | 0.18 | 0.79 | 0.20 | 0.80 | 0.12 | −0.40 | 1.00 | 0.50 |
| a_evap | 0.25 | 0.70 | 0.38 | 0.16 | 0.43 | 0.26 | 0.43 | 0.35 | −0.21 | −0.36 | 0.50 | 1.00 |

**Figure S3:** Pairwise correlation coefficients of the climatic values of native species occurrences for all original 12 climatic variables: diffuse radiation (dif_rad, W/m²), leaf area index (leaf_ai), relative humidity (rel_humidity, %), precipitation seasonality (seasonality, standard deviation of monthly precipitation values), mean maximum temperature of the hottest four months (hot_mean; °C), mean precipitation of the driest three months (dry_avg; mm), mean minimum temperature of the coldest four months (cold_mean; °C), mean precipitation of the wettest three months (wet_avg; mm), soil evaporation (s_evap, mm), solar radiation (s_rad, W/m²), available soil moisture (s_mois), and actual evapotranspiration (a_evap, mm).

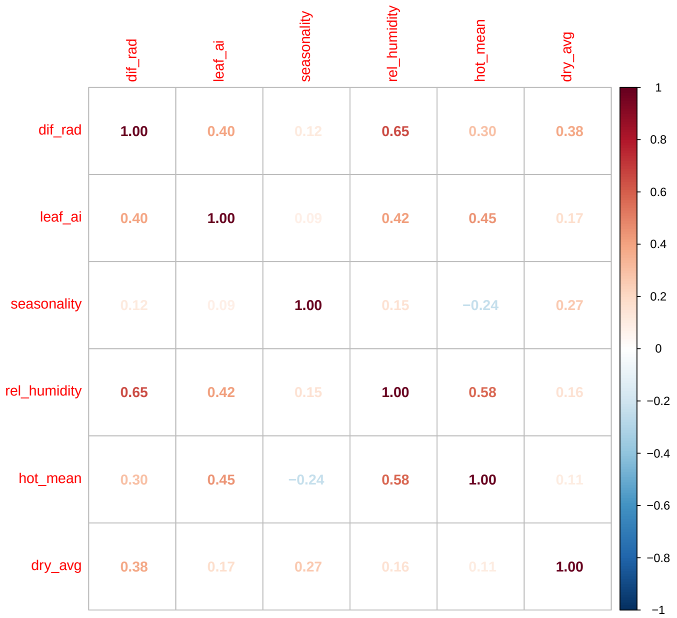


**Figure S4:** Pairwise correlation coefficients of the climatic values of native species occurrences for the six selected climatic variables: : diffuse radiation (dif_rad, $W/m$), leaf area index (leaf_ai), precipitation seasonality (seasonality, standard deviation of monthly precipitation values), relative humidity (rel_humidity, %), mean maximum temperature of the hottest four months (hot_mean; $^{\circ}C$), and mean precipitation of the driest three months (dry_avg; mm).

### Ecological niche models: parameter fit results and suitability maps

Under the approach proposed by Jiménez and Soberón (2022), the underlying model for the fundamental niche is a multivariate normal distribution and the parameters of interest are $\mu$ (the optimal climatic conditions for the survival of the species) and $\Sigma$ (a variance-covariance matrix that determines the shape and size of the niche). We coded the corresponding log-likelihood functions and minimized its negative to obtain the maximum likelihood estimates (MLEs) of these parameters, $\hat{\mu}$ and $\hat{\Sigma}$ (see the functions included in "00_Niche_functions.r" and the code in "04_Niche_modeling_3D.R"). Table S1 contains the MLEs obtained for each model and species. We used these MLEs to plot the 95% confidence ellipsoids under Model 1 and Model 2, and for each species. The resulting ellipsoids are plotted in Figures S5 and S6; they are considered the border of the fundamental niche of each species. In these figures, we added the occurrence points used to fit the models and the occurrence points of the non-native species. In both models, we observe that most of the occurrence of non-native species are inside some of the ellipsoids.

The MLEs were then used to create suitability maps in which the color of a pixel indicates how far the climatic conditions at this site are from the center of the estimate ellipsoid. The resulting suitability maps are shown in Figure S7, where each color represents a different species. The R code used to create these maps can be found in the files "00_Niche_functions.r" and "05_Suitability_maps.R" at the GitHub repository; the latter also includes the code used to binarize the maps and combine the outputs of Model 1 and Model 2.

We performed the whole model fitting procedure using both 8000 and 10000 random points within the Hawaiian islands. In the model, this set of random points is used to calculate weights (ranging from 0 to 1) by which each occurrence records is multiplied. A weight close to zero means that the climatic conditions observed at the corresponding occurrence are rare in the accessible area and, therefore, the probability of observing an occurrence at those climatic conditions should be low. On the other hand, a weight close to one means that the climatic conditions observed at the corresponding occurrence are common in the accessible area and the probability of observing an occurrence at those climatic conditions is high because those conditions are abundant. These weights are calculated by approximating the distribution of climatic conditions within the accessible area through a kernel density estimation, therefore, the larger the number of random points taken from the accessible area the better.

The results between the models fitted with 8000 and 10000 points in the study area are very similar. Therefore, we include only those results obtained with the model fitted with 10000 points. However, we here include maps that shows the differences between these two models, see Figure S8 which shows the differences between the suitability areas predicted with the model that used 8000 points and suitable areas predicted when increasing the sample size to 10000. The R code used to identify the differences in model fit and create these maps is included in the file called “07_Model_comparison.r”

**Table S1:** Maximum likelihood estimates of the parameters that determine the fundamental niche of the four native ferns under Model 1 and Model 2. $\hat{\mu}$ represents the center of the ellipsoidal niche and the matrix $\hat{\Sigma}$ defines the shape and size of the ellipsoids.

| | Model 1 | | Model 2 | |
|---|---|---|---|---|
| Species | $\hat{\mu}$ | $\hat{\Sigma}$ | $\hat{\mu}$ | $\hat{\Sigma}$ |
| *D. angelica* | (91.55, 86.36, 61.40) | $\begin{pmatrix} 13.55 & 4.52 & 39.13 \\ 4.52 & 1.64 & 13.21 \\ 39.13 & 13.21 & 122.82 \end{pmatrix}$ | (3.05, 23.72, 190.53) | $\begin{pmatrix} 0.45 & -0.09 & 2.86 \\ -0.09 & 0.56 & -8.70 \\ 2.86 & -8.70 & 223.84 \end{pmatrix}$ |
| *D. decora* | (88.47, 82.45, 55.83) | $\begin{pmatrix} 35.24 & 8.79 & 22.00 \\ 8.79 & 39.33 & 71.43 \\ 22.00 & 71.43 & 417.85 \end{pmatrix}$ | (1.77, 27.09, 158.59) | $\begin{pmatrix} 2.11 & -1.45 & 50.92 \\ -1.45 & 3.18 & -73.01 \\ 50.92 & -73.01 & 4065.62 \end{pmatrix}$ |
| *D. decipiens* | (87.03, 84.43, 45.56) | $\begin{pmatrix} 130.26 & 104.29 & 78.88 \\ 104.29 & 126.25 & 75.00 \\ 78.88 & 75.00 & 137.55 \end{pmatrix}$ | (2.73, 25.26, 181.32) | $\begin{pmatrix} 1.68 & -2.43 & 55.55 \\ -2.43 & 3.94 & -85.22 \\ 55.55 & -109.18 & 2874.30 \end{pmatrix}$ |
| *P. ternifolia* | (83.82, 65.90, 48.07) | $\begin{pmatrix} 135.02 & 190.85 & 280.50 \\ 190.85 & 448.55 & 828.69 \\ 280.50 & 828.69 & 3053.70 \end{pmatrix}$ | (0.44, 5.54, 208.96) | $\begin{pmatrix} 0.77 & 0.51 & 20.52 \\ 0.51 & 48.21 & -29.22 \\ 20.52 & -29.22 & 4139.40 \end{pmatrix}$ |

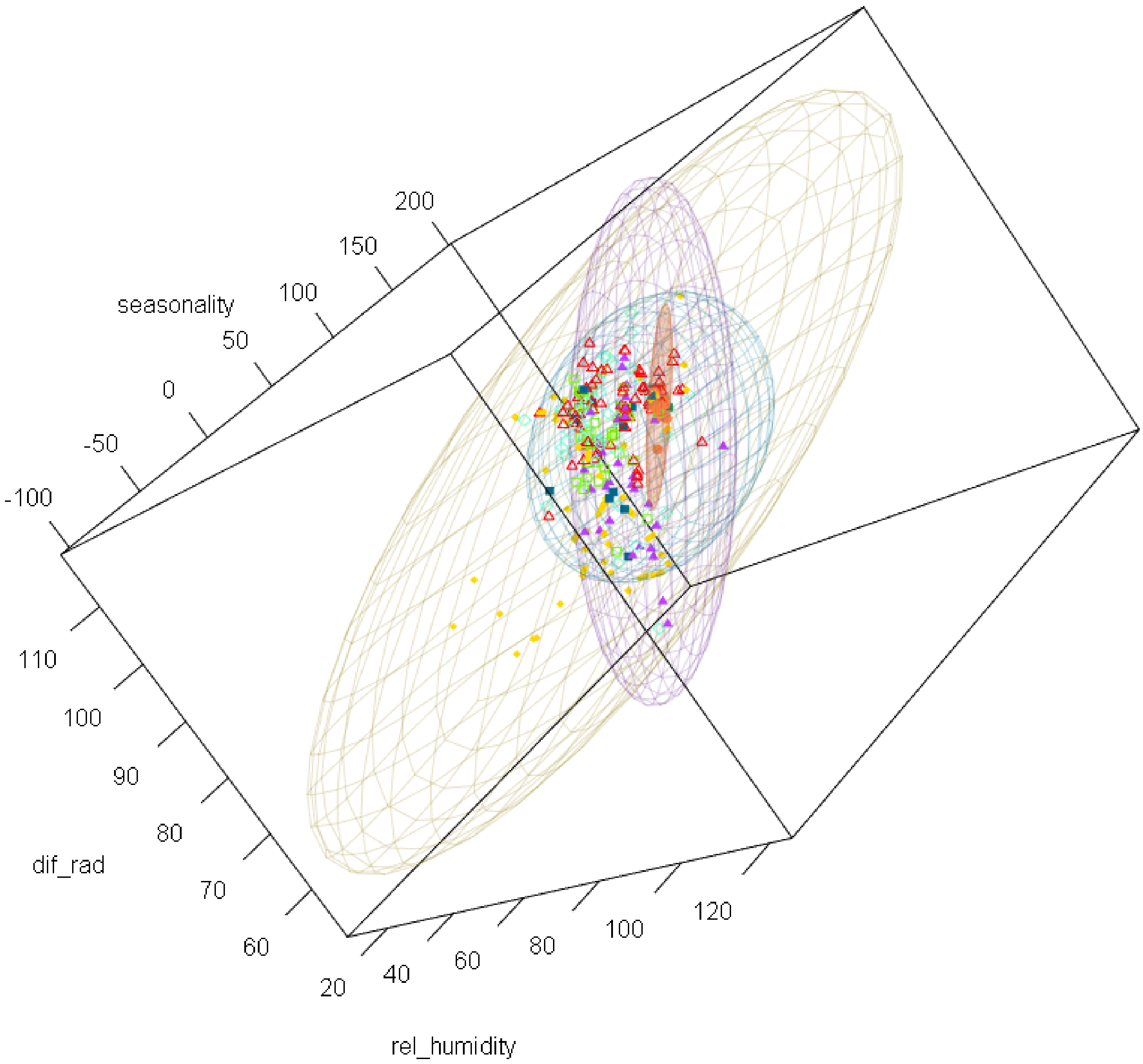


**Figure S5:** Occurrence data for native and non-native dryland ferns in 3D space for Model 1. The three variables used in Model 1 correspond to the three axes in the graph: relative humidity, seasonality (of temperature), and differential radiation. Ellipses correspond to the inferred niche for native species. Non-native species occurrences are shown with points but have no corresponding ellipse.

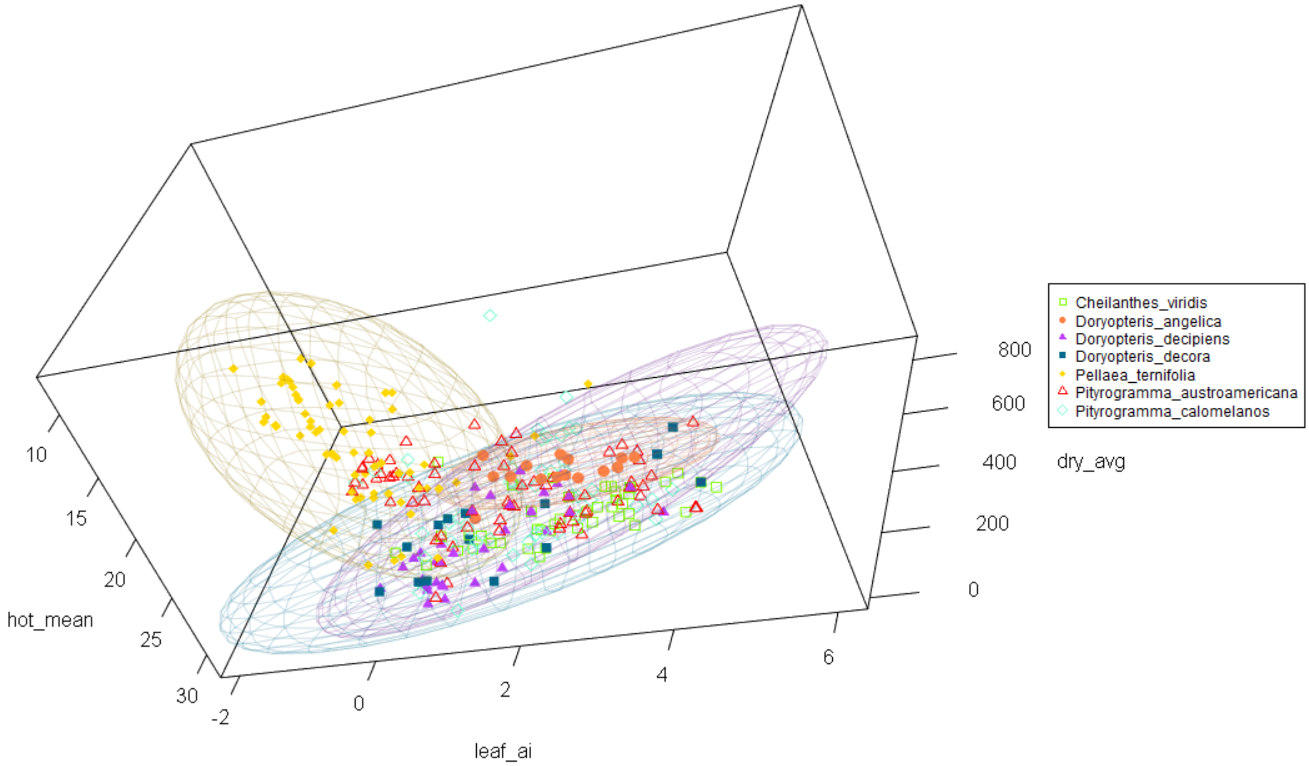


**Figure S6:** Occurrence data for native and non-native dryland ferns in 3D space for Model 2. The three variables used in Model 2 correspond to the three axes in the graph: mean maximum temperature of the hottest season, leaf area index, and the mean minimum precipitation of the driest season. Non-native species occurrences are shown with points but have no corresponding ellipse.

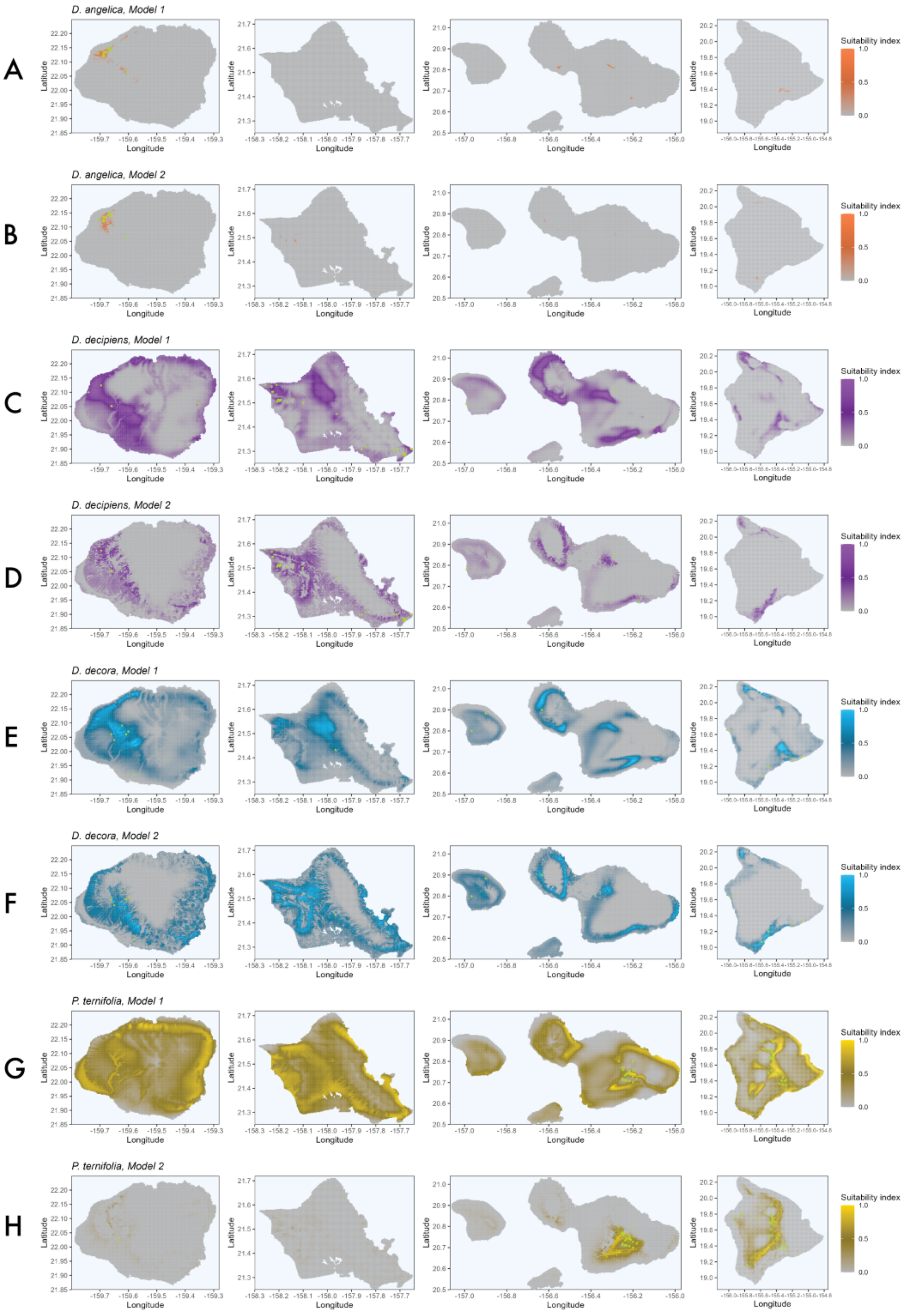


**Figure S7:** Suitability maps obtained with Model 1 and Model 2, respectively. The occurrence points used to estimate the niches for native Hawaiian dryland ferns were drawn in green. (A) and (B) *Doryopteris angelica*, (C) and (D) *Doryopteris decipiens*, (E) and (F) *Doryopteris decora*, and (G) and (H) *Pellaea ternifolia*.

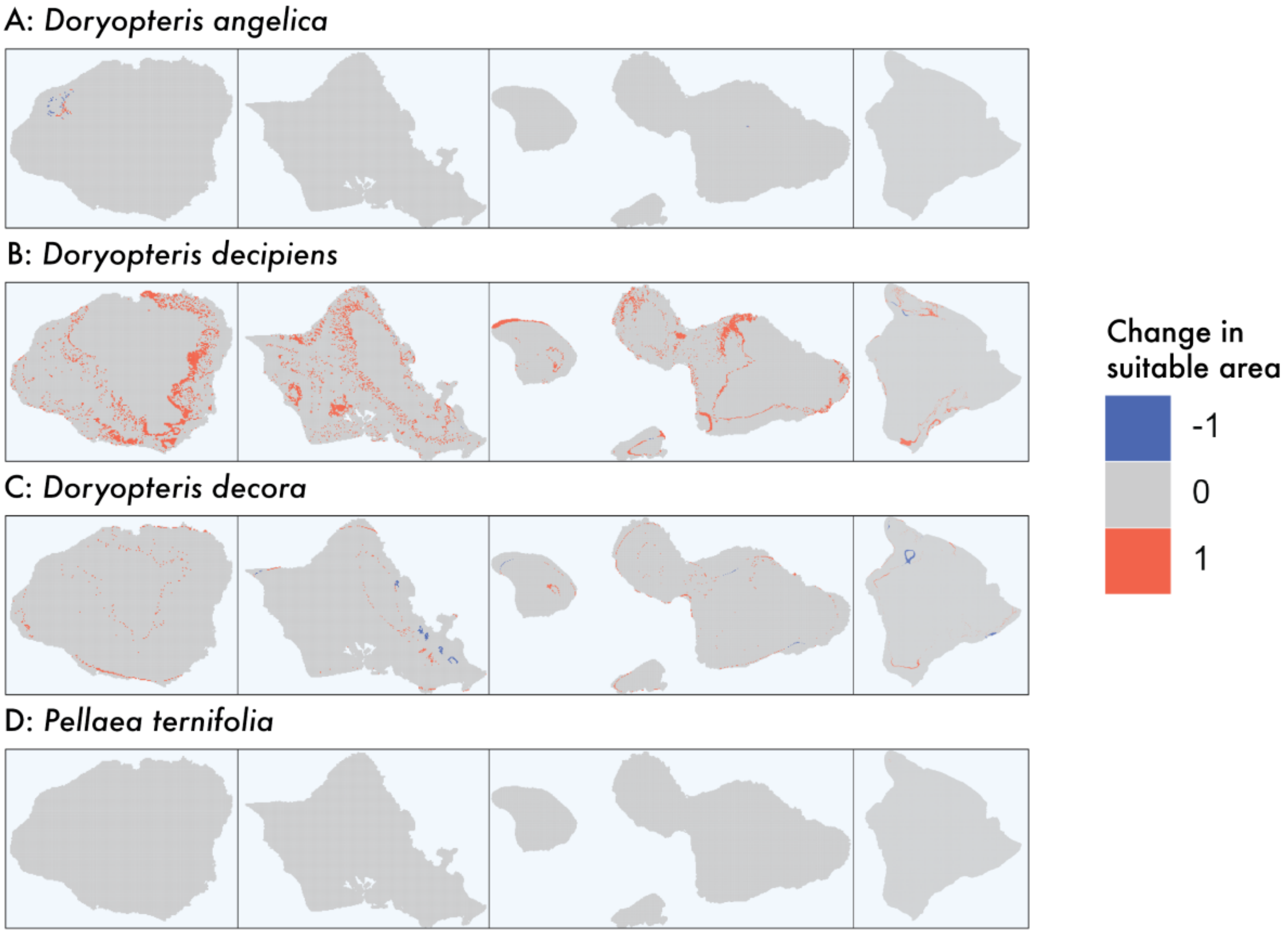


**Figure S8:** Comparison of model outputs: difference between models fitted with 8000 and 10,000 random points to estimate the weights by which each occurrence point is multiplied by in the likelihood function. The areas in blue are predicted as suitable under the model fitted with 8000 points, areas in red are predicted as suitable under the model fitted with 10,0000 points, and areas in grey represent cells were model predictions match. (A) *Doryopteris angelica*, (B) *Doryopteris decipiens*, (C) *Doryopteris decora*, and (D) *Pellaea ternifolia*.

#### Overlap of naturalized species occurrences with native species suitable areas

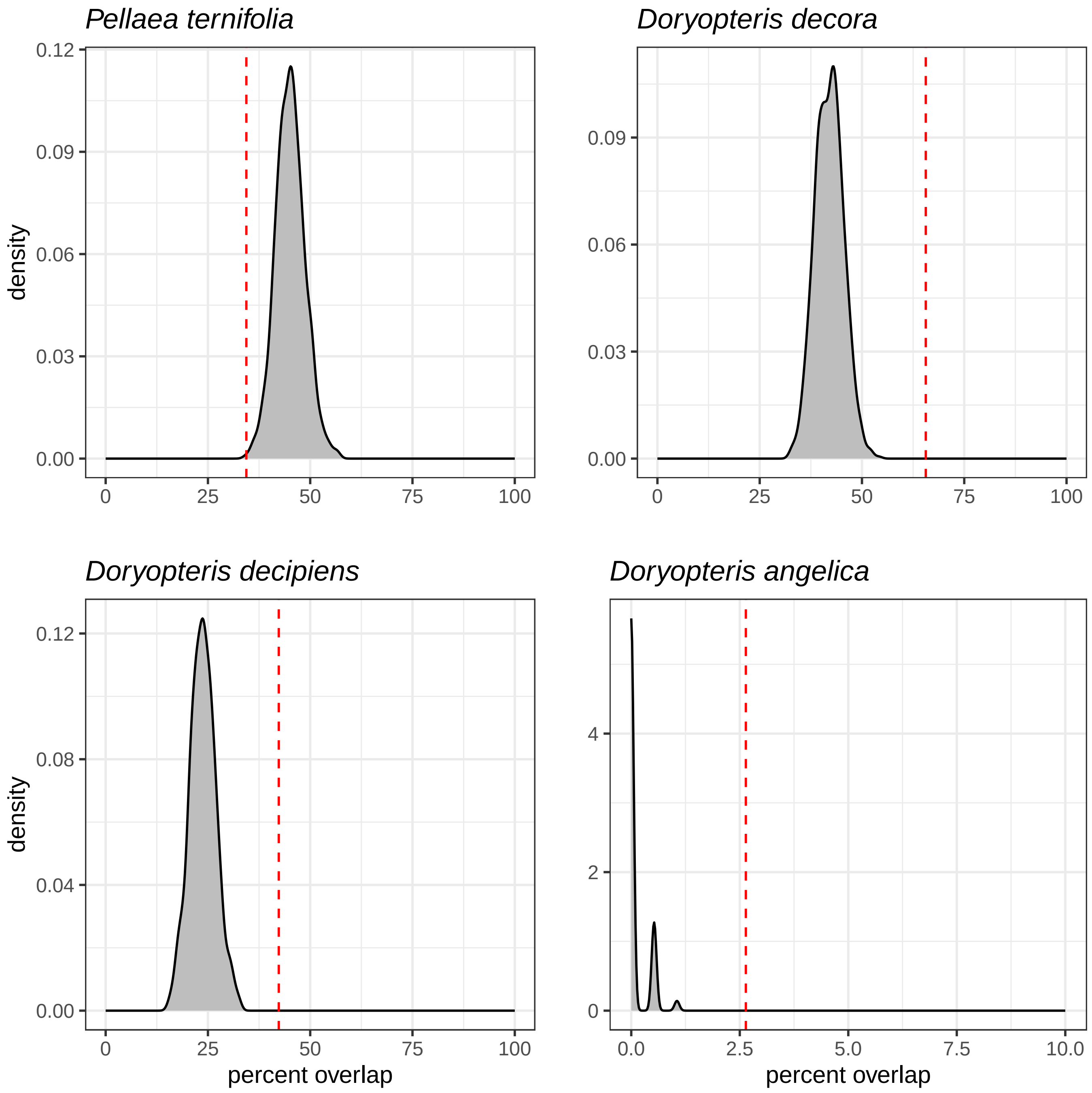


**Figure S9:** Distributions (in grey) of the expected overlap between all naturalized occurrences and the suitable area for each native species. The dashed line (in red) represents the empirical estimate (also labeled in text). When the red line falls outside of the grey distribution, our empirical estimate is significantly different that expected by chance.

#### Land-cover unit reclassification codes and their percentage of intersection with climatically suitable areas

We worked with a raster layer that depicts the land cover and degree of human disturbance to plant communities on the seven main Hawaiian Islands, as downloaded from https://www.sciencebase.gov/catalog/item/592dee56e4b092b266efeb6b (Jacobi et al., 2017). The original 48 units were first converted into 18 new classes based on biomes and general land-cover units. We applied a second reclassification where we considered four habitat statuses: native, mixed (native-alien), alien, and developed, which refer to the degree of disturbance to vegetation. Table S3 displays the old and new labels of the land-cover units.

We used the new land-cover layers and intersected them with the binary maps of models 1 and 2 to calculate the percentage of suitable area covered by each land-cover unit, under both Classification 1 and Classification 2. The resulting percentages are arranged in Table S2. We used the R code included in the file "06_LandCover_Hawaii.R" to create Tables S3 and S2, as well as Figure 5 in the main text.

**Table S2:** Percentage of climatically suitable area covered by each land-cover type included in Classification 1 and Classification 2..

| **Classification 1** | *D. angelica* | *D. decora* | *D. decipiens* | *P. ternifolia* |
|---|---|---|---|---|
| Wet forest | 0 | 5.29 | 2.27 | 6.77 |
| Mesic forest | **80.37** | 15.25 | **11.22** | 7.94 |
| Dry forest | 0 | 7.52 | 12.80 | 6.62 |
| Wet shrubland | 0.31 | 1.02 | 0.25 | 1.11 |
| Mesic shrubland | 7.67 | 4.90 | 4.15 | 8.37 |
| Dry shrubland | 1.53 | **11.10** | 17.36 | 13.82 |
| Wet grassland | 0 | 0.50 | 0.20 | 0.77 |
| Mesic grassland | 0.61 | 7.83 | 6.63 | 10.16 |
| Dry grassland | 0 | 13.43 | 15.04 | 12.86 |
| Alien tree plantation | 4.60 | 1.76 | 1.80 | 0.34 |
| Agriculture | 0 | 7.42 | 7.74 | 1.46 |
| Low intensity developed | 2.15 | 5.43 | 6.01 | 1.30 |
| Medium intensity developed | 0 | 2.15 | 2.87 | 0.52 |
| High intensity developed | 0 | 1.80 | 2.61 | 0.25 |
| Developed open space | 0 | 1.91 | 1.79 | 0.20 |
| Not vegetated | 2.76 | 12.34 | 6.86 | **27.41** |
| Coastal strand | 0 | 0 | 0 | 0 |
| Wetland | 0 | 0.33 | 0.40 | 0.09 |
| **Classification 2** | *D. angelica* | *D. decora* | *D. decipiens* | *P. ternifolia* |
| Native | 59.51 | 11.20 | 7.39 | 34.91 |
| Mixed | 2.76 | 13.56 | 8.23 | 27.77 |
| Alien | 35.58 | 63.94 | 71.10 | 35.05 |
| Developed | 2.15 | 11.29 | 13.28 | 2.27 |

**Table S3:** Reclassification of land-cover units. The original 48 detailed land-cover units (first and second column) were converted into two new classifications: 'Classification 1' combines biome units and detailed development-related units, 'Classification 2' only include four categories that represent the degree of disturbance to vegetation, from 'native' to 'developed'.

| ID | Detailed land-cover unit | Classification 1 | Classification 2 |
|---|---|---|---|
| 1 | Closed ʻōhiʻa wet forest | Wet forest | Native |
| 2 | Closed koa-ʻōhiʻa wet forest | Wet forest | Native |
| 3 | Low-stature ʻōhiʻa wet forest | Wet forest | Native |
| 4 | Open ʻōhiʻa wet forest | Wet forest | Native |
| 5 | Open koa-ʻōhiʻa wet forest | Wet forest | Native |
| 6 | Closed ʻōhiʻa mesic forest | Mesic forest | Native |
| 7 | Closed hala forest | Mesic forest | Native |
| 8 | Closed koa-ʻōhiʻa mesic forest | Mesic forest | Native |
| 9 | Open ʻōhiʻa mesic forest | Mesic forest | Native |
| 10 | Open koa-ʻōhiʻa mesic forest | Mesic forest | Native |
| 11 | Closed ʻōhiʻa dry forest | Dry forest | Native |
| 12 | Mixed mamane-naio-native trees dry woodland | Dry forest | Native |
| 13 | Open ʻōhiʻa dry forest | Dry forest | Native |
| 14 | Open koa-mamane dry forest | Dry forest | Native |
| 15 | Native mesic to dry forest and shrubland | Mesic forest | Native |
| 16 | Mixed native-alien wet forest | Wet forest | Mixed |
| 17 | Mixed native-alien mesic forest | Mesic forest | Mixed |
| 18 | Mixed native-alien dry forest | Dry forest | Mixed |
| 19 | Alien wet forest | Wet forest | Alien |
| 20 | Alien mesic forest | Mesic forest | Alien |
| 21 | Alien dry forest | Dry forest | Alien |
| 22 | Kiawe dry forest and shrubland | Dry forest | Alien |
| 23 | Plantation forest | Alien tree plantation | Alien |
| 24 | Native wet cliff community | Wet shrubland | Native |
| 25 | Native wet shrubland | Wet shrubland | Native |
| 26 | Native mesic shrubland | Mesic shrubland | Native |
| 27 | Mixed native-alien dry cliff community | Dry shrubland | Native |
| 28 | Native dry shrubland | Dry shrubland | Native |
| 29 | Coastal strand vegetation | Coastal strand | Native |
| 30 | Uluhe ferns and native shrubs | Wet shrubland | Native |
| 31 | Mixed native-alien wet shurbs and grass/sedges | Wet shrubland | Mixed |
| 32 | Mixed native-alien mesic shrubs and grass | Mesic shrubland | Mixed |
| 33 | Alien wet shrubland | Wet shrubland | Alien |
| 34 | Alien mesic shrubland | Mesic shrubland | Alien |
| 35 | Alien dry shrubland | Dry shrubland | Alien |
| 36 | Native Deschampsia grassland | Mesic grassland | Native |
| 37 | Alien wet grassland | Wet grassland | Alien |
| 38 | Alien mesic grassland | Mesic grassland | Alien |
| 39 | Alien dry grassland | Dry grassland | Alien |
| 40 | Native bog community | Wetland | Mixed |
| 41 | Wetland vegetation | Wetland | Mixed |
| 42 | Cultivated agriculture | Agriculture | Alien |
| 43 | Developed open space | Developed open space | Developed |
| 44 | High intensity developed | High intensity developed | Developed |
| 45 | Low intensity developed | Low intensity developed | Developed |
| 46 | Medium intensity developed | Medium intensity developed | Developed |
| 47 | Very sparse vegetation to unvegetated | Not vegetated | Mixed |
| 48 | Water | Wetland | Mixed |
